# Assessing the global prevalence of wild birds in trade

**DOI:** 10.1101/2023.08.09.552606

**Authors:** Paul F. Donald, Eresha Fernando, Lauren Brown, Michela Busana, Stuart H.M. Butchart, Serene Chng, Alicia de la Colina, Juliana Machado Ferreira, Anuj Jain, Victoria R. Jones, Rocio Lapido, Kelly Malsch, Amy McDougall, Colum Muccio, Dao Nguyen, Willow Outhwaite, Silviu O. Petrovan, Ciara Stafford, William J. Sutherland, Oliver Tallowin, Roger Safford

## Abstract

Trade represents a significant threat to many wild species and is often clandestine and poorly monitored. Information on which species are most prevalent in trade, and potentially threatened by it, therefore remains fragmentary. We mobilised seven global datasets on birds in trade to identify the species or groups of species that might be at particular risk. These datasets sample different parts of the broad trade spectrum but we nevertheless find a statistically strong congruence between them in which species are recorded in trade. Furthermore, the frequency with which species are recorded within datasets is positively correlated with their occurrence across datasets. This allows us to propose a trade prevalence score that can be applied to all bird species globally. This score discriminates well between species known from semi-independent assessments to be heavily or unsustainably traded and all other species. Globally, 45.1% of all bird species, and 36.7% of globally threatened bird species, were recorded in at least one of the seven datasets. Species listed in Appendices I or II of CITES, species with large geographical distributions and non-songbirds had higher trade prevalence scores. Speciose orders with high mean trade prevalence scores include the Falconiformes, Psittaciformes, Accipitriformes, Anseriformes, Bucerotiformes and Strigiformes. Despite their low mean prevalence score, Passeriformes accounted for the highest overall number of traded species of any order but had low representation in CITES Appendices. Geographical hotspots where large numbers of traded species co-occur differed between songbirds (South-East Asia and Eurasia) and non-songbirds (central South America, sub-Saharan Africa and India). This first attempt to quantify and map the relative prevalence in trade of all bird species globally can be used to identify species and groups of species which may be at particular risk of harm from trade and can inform conservation and policy interventions to reduce its adverse impacts.

**Article impact statement:** The first metric to estimate the prevalence in trade of all the world’s bird species is presented.

## INTRODUCTION

Wild birds are used by people for many purposes, including food, clothing, ornamentation, religious practices, sport and pets. The trade that supplies this demand is recognised as a serious direct threat to the survival of many species (e.g. Tingley *et al*. 2017, IPBES 2019, Ribeiro *et al*. 2019, Cardoso *et al*. 2021, Morton *et al*. 2021). Trade can also pose wider threats through the spread of zoonotic and other infectious wildlife diseases (Gómez & Aguirre 2008, Borsky *et al*. 2020, Borzée *et al*. 2020, Bezerra-Santos *et al*. 2021, Petrovan *et al*. 2021, Rush *et al*. 2021), the establishment of invasive populations (Carrete & Tella 2008, Abellán *et al*. 2016, Souviron-Priego *et al*. 2018), the degradation of community structure and ecosystem services (Española *et al*. 2013, Marthy & Farine 2018), and the impacts of bycatch on non-target species (Khelifa *et al*. 2017). There are also significant animal welfare issues in the trade of wild species (Baker *et al*. 2013). However, trade in wild birds is often unregulated, illegal, clandestine and poorly monitored, so information on which species are most prevalent in trade, and therefore potentially at greatest risk, remains fragmentary and biased.

Previous analyses have suggested that up to 45% of all extant wild bird species are traded to some extent and that the most prevalent form of trade is capture for the pet industry, which might affect as many as 37% of all species (Butchart 2008, BirdLife International 2022). A recent analysis suggests that over 200 globally threatened bird species owe their poor global conservation status at least partly to trade (Challender *et al*. 2023). Many recently published articles on the threats posed by trade to wild birds focus on South-East Asia, where many songbird species are in high demand for the cage-bird industry and the Buddhist practice of ‘merit release’ (e.g. Nijman 2010, Gilbert *et al*. 2012, Lee *et al*. 2016, Harris *et al*. 2017, Chng *et al*. 2018, Rentschlar *et al*. 2018, Marshall *et al*. 2020). Some species in the region have become threatened with extinction by trade (Eaton *et al*. 2015, Shepherd *et al*. 2016, Rheindt *et al*. 2019), and indeed one heavily traded species, Javan Pied Starling *Gracupica jalla*, may now be extinct in the wild (Nijman *et al*. 2021, van Balen & Collar 2021). The exceptional concern this trade has generated led to the formation of the IUCN Species Survival Commission (SSC) Asian Songbird Trade Specialist Group (ASTSG) (Shepherd & Cassey 2017). A heavy trade in songbirds for the cage-bird industry has also been reported in parts of South America (Alves *et al*. 2010, Alves *et al*. 2012, Regueira & Bernard 2012, do Nascimento *et al*. 2015, Souto *et al*. 2017, Charity & Ferreira 2020). Further trade sectors that have received particular attention include the taking of parrots for the pet industry (e.g. Weston & Menon 2009, Gastañaga *et al*. 2010, Marsden & Royle 2015, Annorbah *et al*. 2016, Martin 2018, Aloysius *et al*. 2019, Chan *et al*. 2021, Wang *et al*. 2021), the use of body parts of vultures in Africa for traditional medicines (e.g. Williams *et al*. 2014, Buij *et al*. 2015, Ogada *et al*. 2016, Williams *et al*. 2021, Daboné *et al*. 2022), the capture of raptors for falconry (e.g. Ma & Chen 2007, Wyatt 2009, Levin 2011, Shobrak 2015, Stretesky *et al*. 2018) and trade in certain wild-killed birds for culinary use in Europe and the Middle East (TRAFFIC 2008, Eason *et al*. 2015, Brochet *et al*. 2016, Brochet *et al*. 2017, Brochet *et al*. 2019). Other forms of trade are more localised or species-specific, such as the use of dried hummingbird carcasses in love charms in Mexico (Trail 2022), trade in the casques of the Helmeted Hornbill *Rhinoplax vigil* for making ornaments (Collar 2015, Beastall *et al*. 2016, Krishnasamy *et al*. 2016) and trade in the eggs of Maleo *Macrocephalon maleo* and other megapodes for food (Argeloo & Dekker 2009, Froese & Mustari 2019, Summers *et al*. 2023). New forms of trade can arise quickly in response to cultural trends; for example a recent surge in trade in owls in Indonesia might be at least partly associated with the popularity of the Harry Potter books (Shepherd & Shepherd 2009, Nijman & Nekaris 2017, Siriwat *et al*. 2020).

In response to such pressures, most national governments have legislation to limit the most damaging exploitation. In addition, many governments are Parties or Signatories to multilateral environmental agreements, many of which confer specific obligations with regard to ensuring trade is sustainable, legal and safe through monitoring and regulation. For example, for Parties to the Convention on the Conservation of Migratory Species of Wild Animals (CMS), trade in CMS Appendix I species is prohibited other than in some specified exceptional circumstances. The primary international policy instrument to regulate international trade in threatened species is the Convention on International Trade in Endangered Species of Wild Fauna and Flora (CITES), to which 183 of the world’s countries are now Parties. This agreement subjects international trade in selected species to certain controls, including authorisation of the trade through a permitting system that considers both legality and sustainability. The species covered by CITES are listed in three Appendices. Appendix I lists threatened species whose commercial trade is prohibited (subject to certain exemptions and special procedures) and whose non-commercial trade is permitted only in exceptional circumstances. Appendix II lists species whose trade must be controlled in order to avoid utilization incompatible with their survival. Appendix III lists species that are protected in at least one country, which has asked other CITES Parties for assistance in controlling their trade. Currently, CITES lists 156 species of birds in Appendix I, 1,294 species in Appendix II and 60 species in Appendix III. However, the CITES website notes that these numbers are approximate ‘because there are no agreed lists for some of the higher taxa’. Most birds covered by Appendix II are listed at the level of family or order, largely to reduce the problems of identifying traded birds to species level. For example, CITES includes all species of the family Trochilidae (hummingbirds) in Appendix II (with a single exception, Hook-billed Hermit *Glaucis dohrnii*, listed in Appendix I), thereby avoiding differences of taxonomic opinion about how many species of hummingbirds there are in the world.

CITES monitors the legal transactions of the international trade in CITES-listed species, and CITES Parties have recently started submitting information on seizures of CITES-listed species through annual reports on illegal trade. However, trade in the majority of species that are not CITES-listed, particularly illegal trade, is poorly monitored. Illegal trade is by its very nature clandestine, and efforts to monitor it are largely limited to information gleaned from surveys of markets (including online markets) and from occasional seizures of illegal shipments of animals or their body parts. Legal trade in non-CITES listed species is monitored equally patchily, primarily through import records to certain countries or blocs with the capacity to do so (such as the EU and USA). All of these different trade surveillance approaches are likely to suffer from inherent but different biases in the way data are collected and the degree to which they represent the underlying trade flows that they attempt to monitor. For example, live birds and larger species are harder to transport and may be more detectable in illegal trade than dead birds and smaller species, and birds being traded within countries may be less detectable than those in international trade. Poor and potentially biased monitoring of trade and its impacts, particularly with regard to non-CITES species, means that the sustainability or otherwise of legal trade is often no easier to assess than that of illegal trade (Andersson *et al*. 2021, Hughes *et al*. 2023), and that the effectiveness of policy interventions is hard to quantify (Challender *et al*. 2015). This in turn means that genuinely sustainable trade can be difficult to identify and may be censured alongside unsustainable trade (Abensperg-Traun 2009, Cooney & Abensperg-Traun 2013, Challender *et al*. 2019, Roe *et al*. 2020).

Because of this incomplete and biased monitoring, there is currently no standardised assessment of prevalence in trade for all the world’s bird species that enables the identification of taxonomic or geographical hotspots of trade. Here, we develop a metric of prevalence in trade by combining a number of disparate global datasets and evaluating their consistency in the frequency and numbers in which species are reported. In doing so, we build on and expand the work of Juergens *et al*. (2021), who compiled comparable data for songbirds. We assess geographic and taxonomic patterns of concordance between datasets to derive a simple score of prevalence in trade that can be used to help identify species that might be heavily traded, and regions where large numbers of traded species co-occur and hence where trade may be of particular concern.

## METHODS

### Data collection and collation

#### Global datasets

We brought together the six largest global datasets that can inform assessment of trade in wild birds. Additionally, we created a new independent dataset by extracting information from published and unpublished reports of birds in trade, largely in physical or online markets; these data were not duplicated in any of the global datasets. We checked for and removed duplication of records within and between datasets, retaining records in the dataset we considered most likely to have first incorporated them. A description of each dataset, and of how we filtered the data for use in each case, is given in Table 1. Six of the datasets provided information on the frequency with which species were recorded in trade and for five of these we had data on the overall numbers of individuals of each species involved. However, sample sizes were generally lower for the latter because not all recording events (e.g. seizures or licencing events) had data on the number of individuals or specimens involved, and some were reported in units other than individual birds (e.g. feathers or bones). We aligned all the datasets to the taxonomy of Handbook of the Birds of the World & BirdLife International (2021), as used by the IUCN Red List, to allow them to be directly compared and combined. This alignment is unlikely to have been perfect, since (a) it was not always possible to determine which taxonomic concept was referred to in each dataset and (b) different taxonomic concepts may have been referred to under the same scientific name, and vice versa, within and between datasets. However, the proportion of records that did not match directly was low, and a high proportion of those will have been aligned correctly, so any errors are unlikely to affect our conclusions materially. We did not include data from the Songbirds in Trade Database (Juergens *et al*. 2021) because it includes data only on Passeriformes and because its data sources include some of the global datasets we analysed.

**Table 1.** Description of the datasets used in the analyses. A general filter was applied across all quantitative sets: for estimating the numbers of birds, we filtered out all entries whose counts are unlikely to relate to whole birds (e.g. those listed as feathers, bone pieces, skin pieces etc.).

| Abbreviated name | Full name (compiler/manager) | Type of trade covered | Species covered | Years covered | Filters applied | No. species recorded | Summary of data included in analyses |
| --- | --- | --- | --- | --- | --- | --- | --- |
| WiTIS | Wildlife in Trade Information System (TRAFFIC)<br>https://www.wildlifetradeportal.org/ | Domestic and international; illegal | Any | 2005-2019 | None | 736 | Quantitative: Data on frequency (number of seizures in which species is reported) and total number of individuals of each species recorded across 2,217 seizures |
| CITES | CITES Trade Database (UNEP-WCMC on behalf of the CITES Secretariat)<br>https://trade.cites.org/ | International; legal | All species included in the CITES Appendices and Annex D of the EU Wildlife Trade Regulations. Includes some species that have been de-listed and so are | 2005-2021 | Direct and indirect trade; all purpose codes; source = wild, unknown or blank; units = blank or number of specimens; terms = whole organism equivalents, defined as live, bodies, | 892 | Quantitative: Data on frequency (number of transactions species reported in) and total number of individuals of each species recorded in international trade. Number of transactions is based on the higher of exporter- or |
|  |  |  | no longer listed in CITES Appendices |  | eggs, eggs (live), skeletons, skins, trophies and skulls |  | importer-reported data. Number of individuals is based on a calculation of gross exports (i.e. the number of individuals traded according to exporters and importers was compared, and the highest value taken). |
| Market surveys | Digitised dataset of market survey reports and other sources of information on birds in trade (BirdLife International, compiled for current study) | Domestic and international; legal and illegal (in unknown proportions) | Any; mostly (>77%) species native to country of market | 2001-2022 | None; data were checked for independence from WiTIS seizure data | 2,008 | Quantitative: Data on frequency (number of surveys species reported in) and total number of individuals of each species recorded across 97 published or unpublished surveys of birds in trade, mostly market surveys (see Table S3) |
| Red List | IUCN Red List data managed in the Species Information System (SIS) (BirdLife International/IUCN)<br>https://www.iucnredlist.org/assessment/sis | Domestic and international; legal and illegal (in | All; the usage coding is applied to | n/a | Non-subsistence usage only | 3,592 listed as being in non- | Qualitative coding of types of use for all the world's birds, with scale of |
|  |  | unknown proportions) | all the world's birds |  |  | subsistence usage | use recorded as one or more of International, National, and Local/subsistence. The first two classes were classed as trade. Underlying data come from multiple sources (Butchart 2008) |
| EU TWIX | EU Trade in Wildlife Information Exchange (Belgian Federal Police and TRAFFIC)<br>https://www.eu-twix.org/ | International (imports to the EU); illegal | Any, but focus on CITES-listed species | 2005-2019 | Excludes seizures of birds known to be of captive origin | 664 | Quantitative; Data on frequency (number of seizures species reported in) and total number of individuals of each species recorded across 5,265 seizure events of birds illegally imported into the EU |
| LEMIS | United States Fish and Wildlife Service's (USFWS) Law Enforcement Management Information System (LEMIS) | International (imports to the USA); mostly legal | Any | 2005-2020 | Excludes animals bred or born in captivity or commercially, hunting trophies, and imports for scientific, biomedical or educational use | 1,556 | Quantitative; Data on frequency (number of events species reported in) and total number of individuals of each species recorded (based on whole-bird metrics) across filtered 10,029 import events to the USA |
| World WISE | The World Wildlife Seizures (World WISE) database (UNODC)<br>https://www.unodc.org/unodc/en/data-and-analysis/wildlife.html | International; illegal | Largely CITES-listed species (60.0%), but non-CITES species also reported when found in seizures of CITES-listed species | 2004-2022 | Data from LEMIS were excluded; wild-caught species only | 1,364 | Quantitative; data on number of seizures involving each species (data on numbers of individual birds and total number of seizures unavailable) |

#### Independent test data

We compiled a list of species identified by field studies or by experts as being heavily or unsustainably traded to assess the level of representation of these species in the seven global trade datasets (Supplementary Tables S1, S2). This list was used as a consistency check of the results generated from the global datasets, and was not intended to be comprehensive. First, we searched for ‘single species’ publications highlighting the threats of trade to individual species or small numbers of allied species that were based on field or other primary data independent of the seven global datasets. We derived these from searches in Google Scholar using the search terms ‘trade’ and ‘bird/s’ until a sample of 50 species was identified (Supplementary Table S1).

Next, we undertook a questionnaire survey of experts in the trade in wild birds to collect information species they considered to be particularly heavily or unsustainably traded within the taxonomic groups or regions of their expertise. Again, this aimed to generate a sample of species known to be heavily or unsustainably traded, not a comprehensive list. We approached potential respondents known to be actively involved in work on the bird trade. Completed questionnaires were provided by representatives of the IUCN Asian Songbird Trade Specialist Group (ASTSG), Asociación Rescate y Conservación de Vida Silvestre (Guatemala), Association Biom (Croatia), Aves Argentinas (Argentina), Burung Indonesia (Indonesia), Freeland Brasil (Brazil), the IUCN Hornbill Specialist Group, One Earth Conservation (Central America), RSPB (UK) and the Temaiken Foundation (Argentina). As questionnaire respondents included some of the authors of studies included in the sample of 50 species, and as those studies would have contributed towards other respondents’ questionnaire responses, there was unsurprisingly a strong overlap between the two lists. Of the sample of 50 species known from published studies to be threatened by trade, 36 (72%) were also included in the questionnaire responses. We therefore combined the two lists to derive a sample of 265 species known or considered from sources independent of the seven trade datasets to be traded to an extent that might affect their conservation status (Supplementary Table S2).

As further semi-independent lists of unsustainably traded bird species, we used the 202 species identified from analyses of Red List data by Challender *et al*. (2023) as likely to be threatened by international trade, and the 27 Tier 1 priority species (comprising 43 taxa) of the Asian Songbird Trade Specialist Group (2022).

### Data analysis

#### Congruence between datasets

We first assessed the extent to which the seven datasets were congruent in which species they recorded in trade, and therefore how meaningfully they could be combined to generate a metric of prevalence in trade for further analysis. We first assigned a value of 1 or 0 to each species in each dataset according to whether or not it was recorded in trade at least once (or in the case of the Red List dataset, whether it was recorded as being in ‘non-subsistence usage’), and then summed these values across datasets to give a value for each species between 0 (species not recorded in any dataset) and 7 (species recorded in all trade datasets). Pearson χ^2^ was used to compare the observed distribution of scores to the frequency of scores that would be expected under the null hypothesis that there was no association between datasets (i.e. that a species recorded in one dataset was no more or less likely to be recorded in another dataset than would be expected by chance). The expected frequency for each species was generated by randomly selecting (with replacement) an observed value from each dataset and summing them to generate a randomised value of between 0 and 7 for each species. We did this for all species globally, for songbirds (Order Passeriformes) and non-songbirds separately, and for species occurring in each of five broad bioregions separately (Americas, comprising Nearctic and Neotropical, Afrotropical, Palearctic, Indomalayan, and Australasian and Oceanian combined), to assess whether there were taxonomic or biogeographical differences in congruence between datasets. We randomly generated expected values from observed values for songbirds and non-songbirds separately, to account for the fact that observed values were not randomly distributed among the two groups of species (see Results).

We then undertook a finer scale assessment of congruence by statistically comparing the observed with the expected frequencies of all 2*^n^* – (*n* + 1) possible two-or-more-way intersections between the *n* = 7 datasets, giving 120 possible intersections. We used the R package *SuperExactTest* (Wang *et al*. 2015) to generate Fisher exact probabilities for all the observed multi-set intersections and to visualise the results. Unlike the previous analysis, this approach is inclusively hierarchical, such that a species recorded in the three-way intersection of datasets *x*, *y* and *z* would also be included in the three two-way intersections of *x* and *y*, *x* and *z,* and *y* and *z*.

We then assessed whether the 0–7 summed score across datasets captured meaningful information on the relative prevalence of species within datasets. We compared the frequency of reporting (the number of times each species was recorded, typically the number of incidents, trade transactions or seizures) and the total number of individual birds (summed across reporting events) of each traded species within each of the six quantitative datasets (see Table 1) with the summed frequency with which that species was recorded (presence/absence) in the other six datasets (a value of 0 to 6). Reporting rates and numbers were logged to normalise the data and ANOVA with *post hoc* Tukey HSD tests used to assess significant pairwise differences in average frequency or abundance between the seven trade prevalence score classes.

We used non-parametric rank-order tests to compare the summed occurrence across the seven trade datasets (a score of 0 to 7) of the 265 species identified in the independent test data (i.e. whose trade has been the subject of one or more single-species publications and/or were listed in the questionnaire survey), with that of all other species combined. We applied the same approach to the list of 202 species identified by Challender *et al*. (2023) as likely to be threatened by international trade, and to the list of 27 priority species of the ASTSG. Because Challender *et al*. used data in the Red List database, we excluded the Red List coding of non-subsistence usage from the combined score for this comparison, which therefore ranged from 0 to 6. In each comparison, our expectation was that if the occurrence of species across the global datasets represents a realistic metric of their prevalence in trade, then those species whose trade has been identified semi-independently as being of particular concern should have a higher median score than all other species and, as a more stringent test, a higher median score than all other species that were recorded at least once in trade.

#### Predictors of prevalence in trade

We compared the prevalence of species in trade (measured as a 0–7 score) between different taxonomic or geographical groupings of species using Wilcoxon rank sum tests with a continuity correction or Kruskal-Wallis tests. In cases where the Kruskal-Wallis test indicated significant between-group differences, we used *post hoc* pairwise Wilcoxon tests to assess where such differences lay between each combination of pairs. We also compared continuous variables such as extent of occurrence (EOO) and generation length between species by log-normalising the data and using ANOVA with *post hoc* Tukey HSD tests.

In order to assess how different explanatory factors interacted in predicting prevalence in trade, we created an ordered three-level factor with nominal levels of “Not traded” (if the species was not recorded in any of the trade datasets), “Low trade score” (for species recorded in one or two datasets) and “High trade score” (for species recorded in three or more datasets). The cut-off point for the second and third groups was selected such that they retained approximately equal numbers of species. We then used ordinal logistic regression with the function *polr* in the *R* package *MASS* (Venables & Ripley 2002, R Core Team 2022) to model this ordered three-level factor as a function of (1) CITES status (a two-level factor of whether a species is listed in CITES Appendices I or II, or not listed), (2) extent of occurrence, derived as the area within a minimum convex polygon around the range map for each species in the dataset of BirdLife International and Handbook of the Birds of the World (2022), (3) generation length (in years) derived from BirdLife’s assessments of extinction risk for all bird species for the IUCN Red List, and (4) a binary songbird *vs* non-songbird factor. Global Red List category was not included in the models because it was strongly correlated with extent of occurrence. Because CITES status was included as a predictor in the models, the contributions of the CITES and World WISE trade datasets to the trade prevalence score were removed when assigning species to the ordinal trade variable, otherwise CITES-listed species would have inflated scores compared to non-CITES species. Extent of occurrence was logged prior to analysis, and the small proportion of species (< 0.2%) with generation lengths over 15 years (all of them non-songbirds) had their generation length truncated to 15 years to avoid extrapolation by the model beyond the range of adequately sampled values. The parallel regression assumption (i.e. that a single set of regression coefficients is sufficient to describe differences across all levels of the ordered trade class factor) was assessed using the graphical method of Harrell (2001).

#### Distribution of trade prevalence

We mapped the distribution of different groupings of species with high trade prevalence scores using a new set of fractional Area of Habitat (AOH) maps at 5-km resolution (ESRI 54017). AOH maps show the parts of each species’ mapped range that fall within its habitat tolerance and altitudinal limits (Brooks *et al*. 2019). Maps were prepared following the protocol of Lumbierres *et al*. (2022b), using the habitat-to-land-cover translation table of Lumbierres *et al*. (2022a), and validated using the protocol of Dahal *et al*. (2022); further details are given in Busana *et al*. (2022). Each pixel value in an AOH map represented the fraction of the pixel suitable for the species; this was converted to a binary value by assigning any pixel with a value larger than 10% to a pixel value of 1 (‘present’) and those with values below this to a value of 0 (‘absent’). For each group, AOH maps were stacked and the number of species whose AOH intersected each 5-km pixel was summed and displayed to highlight regions of the world where large numbers of species that are prevalent in trade co-occur. Both the breeding and the non-breeding distributions of migratory species were included. Pixels with high summed values are those where a large number of traded species co-occur. However, these pixels are not necessarily areas where trade actually takes place, and hotspots of co-occurring traded species are not necessarily the places where trade is most damaging to the survival of species.

We mapped separately the distributions of species with high trade prevalence scores for (1) all species, (2) songbirds (Passeriformes), (3) non-songbirds, (4) CITES-listed species (Appendices I and II) and (5) non-CITES-listed species. The threshold for inclusion differed between groups in order to keep the number of species being mapped roughly comparable between groups, given that average scores differed between them. Thus the threshold for songbirds (species with scores of 3 or above were mapped; n = 423) was lower than that for non-songbirds (species with scores of 4 or above were mapped; n = 444) because songbirds had generally lower scores (see Results). In each case the number of maps included was slightly lower than the number of species in the respective group because a small proportion of species, mostly pelagic seabirds, lacked AOH maps (of the 2174 species with sufficiently high trade prevalence scores to qualify for inclusion in at least one map, 60 species [2.8%] lacked maps).

## RESULTS

Globally, 4,915 species (44.7% of all bird species) were recorded in at least one of the seven trade datasets (Figure 1, Table 2). Of the traded species, 2,290 (46.6%) were recorded in a single dataset, and the remaining species were recorded in two or more (Figure 1).

**Figure 1.**
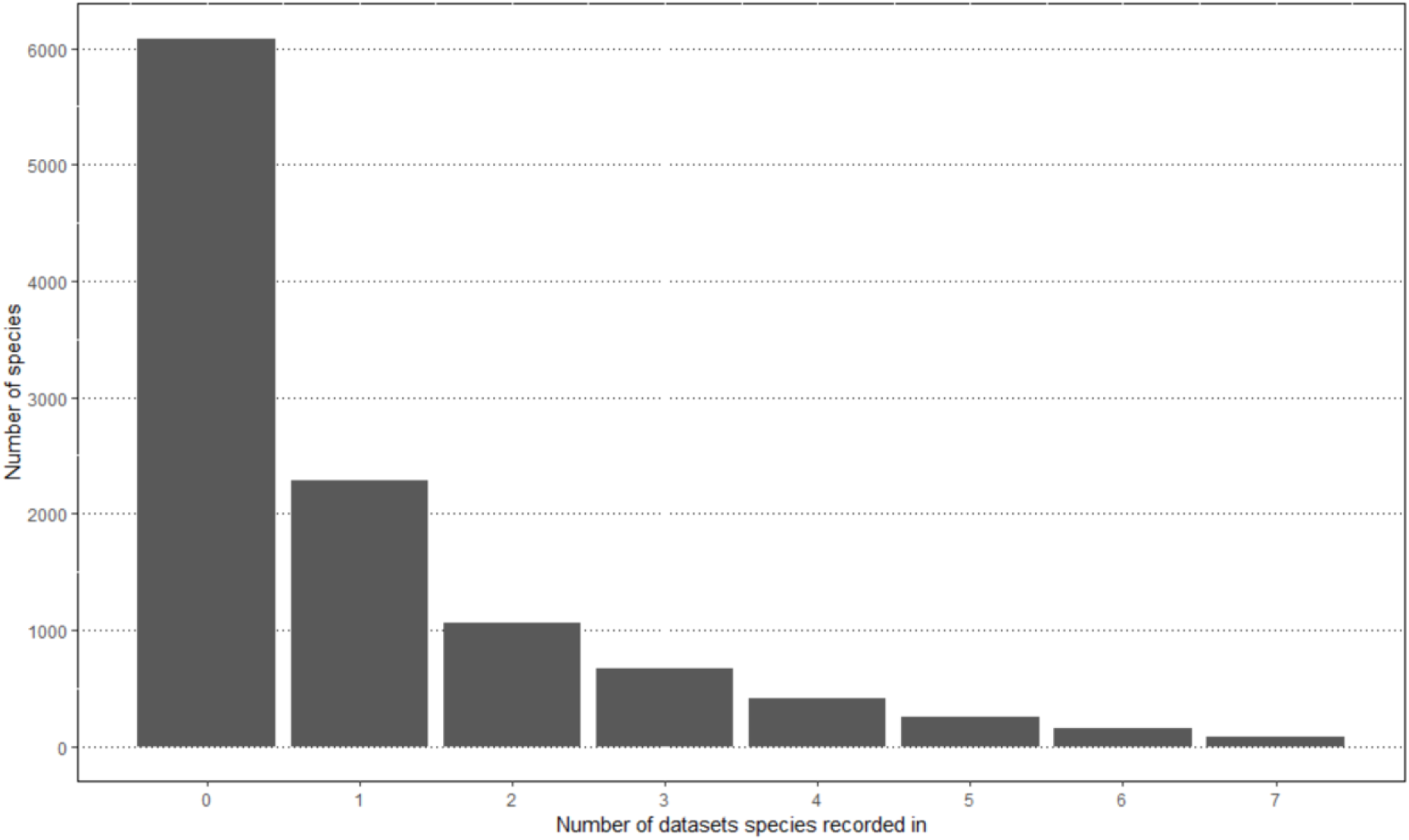
Frequency with which bird species (n = 10,999 globally) were recorded in the seven trade datasets (the trade prevalence score).

**Table 2.** Number of extant species, mean number of datasets (trade prevalence score) per species, number (and %) of species recorded in at least one of the seven trade datasets, number (and %) of species recorded in at least four of the seven trade datasets and number (and %) of species listed in CITES Appendices I or II, for all birds combined and for all orders of birds containing more than 50 species. Orders are listed in descending order of their mean trade prevalence score. Species numbers follow Handbook of the Birds of the World & BirdLife International (2021).

| Order | No. species | Mean trade prevalence score | No. species with score<br>≥1 (%) | No. species with score<br>≥4 (%) | No. species in CITES Appendices I<br>or II (%) |
| --- | --- | --- | --- | --- | --- |
| All birds | 10,999 | 0.98 | 4,915 (44.7) | 896 (8.1) | 1588 (14.4) |
| Falconiformes | 64 | 3.59 | 63 (98.4) | 29 (45.3) | 64 (100) |
| Psittaciformes | 404 | 3.34 | 360 (89.1) | 193 (47.8) | 400 (99.0) |
| Accipitriformes | 251 | 3.09 | 233 (92.8) | 111 (44.2) | 251 (100) |
| Anseriformes | 170 | 2.46 | 155 (91.2) | 36 (21.2) | 15 (8.8) |
| Bucerotiformes | 72 | 2.12 | 54 (75.0) | 14 (19.4) | 29 (40.3) |
| Strigiformes | 238 | 1.81 | 166 (69.7) | 45 (18.9) | 238 (100) |
| Pelecaniformes | 110 | 1.78 | 85 (77.3) | 17 (15.5) | 7 (6.4) |
| Caprimulgiformes | 597 | 1.67 | 354 (59.3) | 123 (20.6) | 366 (61.3) |
| Galliformes | 307 | 1.50 | 196 (63.8) | 44 (14.3) | 35 (11.4) |
| Suliformes | 53 | 1.49 | 50 (94.3) | 1 (1.9) | 2 (3.8) |
| Columbiformes | 353 | 1.07 | 193 (54.7) | 21 (5.9) | 7 (2.0) |
| Charadriiformes | 377 | 0.93 | 201 (53.3) | 7 (1.9) | 4 (1.0) |
| Gruiformes | 169 | 0.86 | 71 (42.0) | 12 (7.1) | 16 (9.5) |
| Coraciiformes | 188 | 0.80 | 78 (41.5) | 8 (4.3) | 0 (0) |
| Piciformes | 484 | 0.64 | 176 (36.4) | 14 (2.9) | 6 (1.2) |
| Cuculiformes | 150 | 0.63 | 59 (39.3) | 1 (0.7) | 0 (0) |
| Passeriformes | 6,603 | 0.58 | 2,236 (33.9) | 184 (2.8) | 87 (1.3) |
| Procellariiformes | 145 | 0.23 | 20 (13.8) | 1 (0.7) | 1 (0.7) |

### Congruence between datasets

When the presence of species recorded in trade was summed across the seven datasets, generating a score of 0–7 for each species, and compared with a score generated by randomly re-assigning values between species within each dataset and summing them, there was a highly significant difference in distribution between classes (Table 3). In each case, significantly more species were recorded in none of the trade datasets or in many datasets than would be expected by chance. This applied to all species globally, to songbirds and non-songbirds separately and when species were broken down by broad bioregion of their occurrence (Table 3).

**Table 3.**
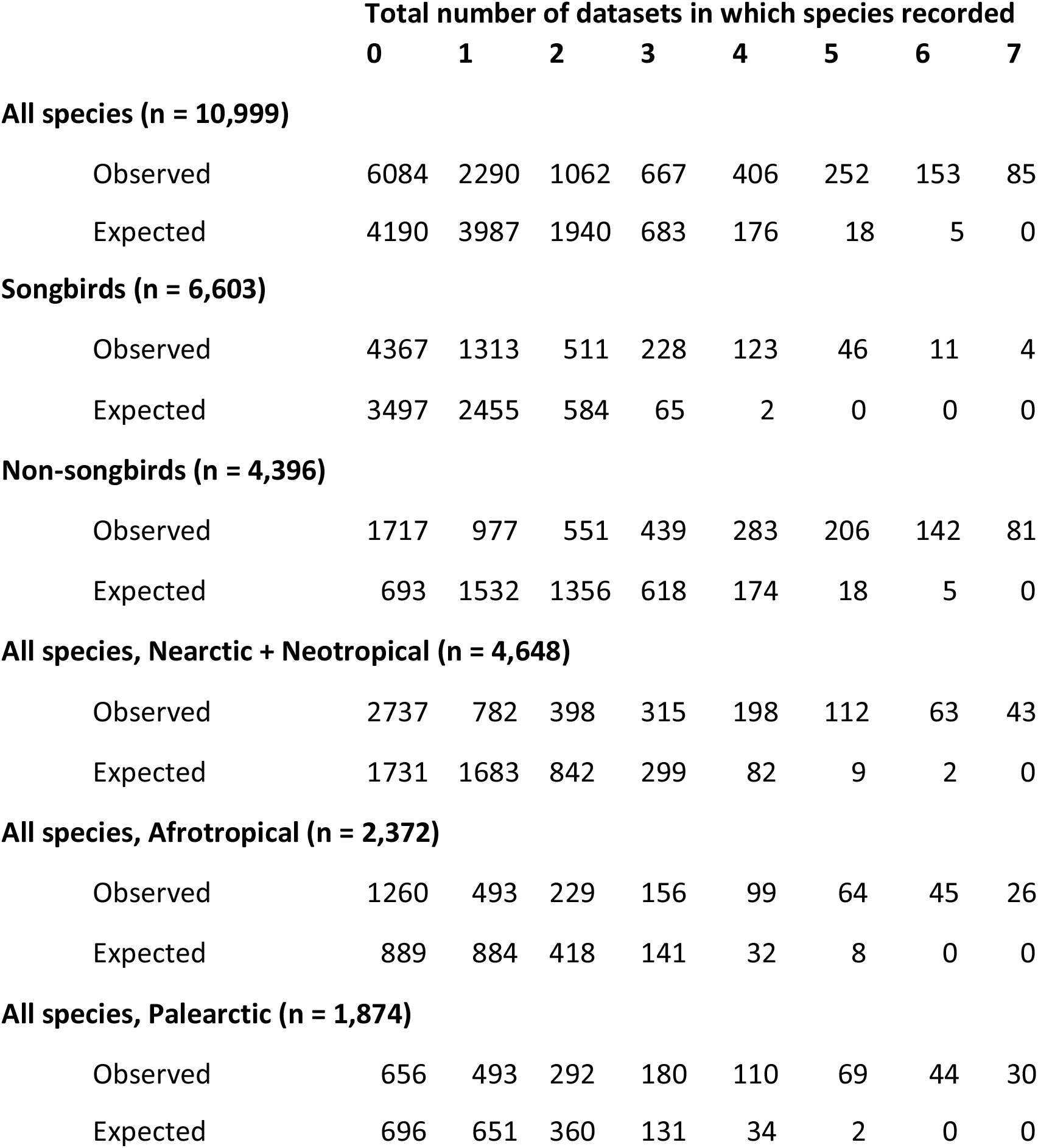
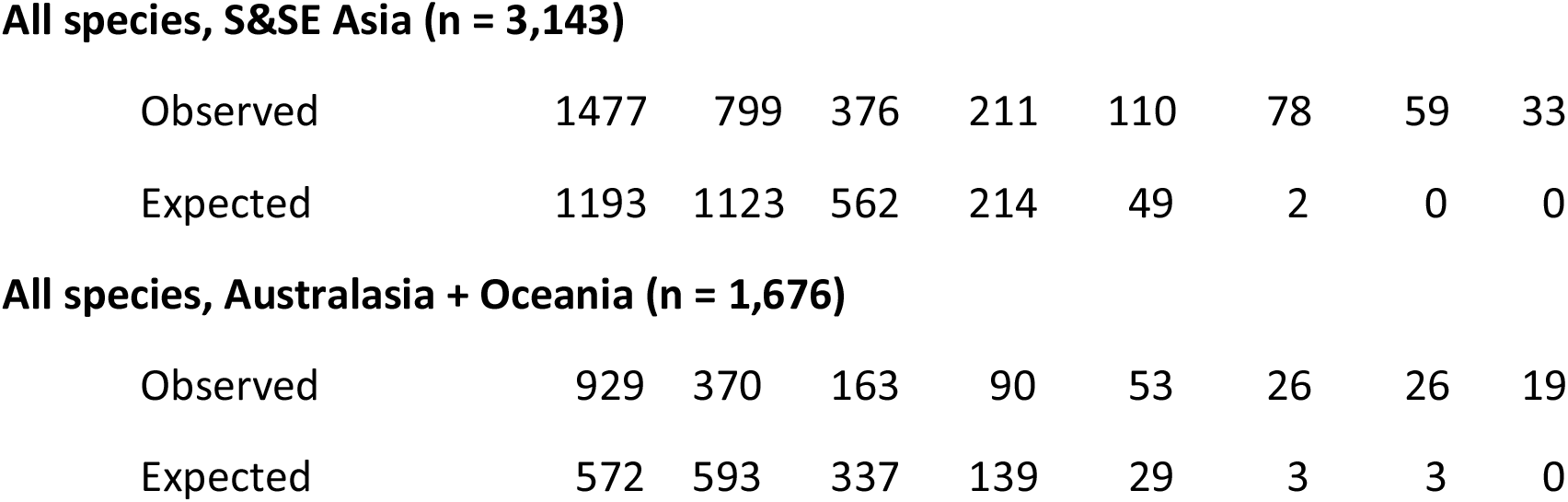
Observed occurrence of 10,999 extant bird species in seven trade datasets and their expected occurrence from the same number of values drawn randomly from the observed data in each dataset. In each case songbirds and non-songbirds were sampled separately to account for the non-random distribution of traded species in each group. There was a highly significant difference between observed and expected totals across all species, for songbirds (passerines) and non-songbirds separately, and for species occupying each bioregion (χ^2^ > 5000, *p* <10^-10^ in all cases), showing that there was a strong tendency for the same species to be recorded in different trade databases that was consistent across regions and taxonomic groupings. Where no species reached an expected score of 5, 6 or 7, observed scores were summed into the highest non-zero randomised score class in the calculation of the Pearson χ^2^. Summed totals across bioregions exceed the count of all species globally because many species occur in two or more bioregions.

|  | Total number of datasets in which species recorded |  |  |  |  |  |  |  |
| --- | --- | --- | --- | --- | --- | --- | --- | --- |
|  | 0 | 1 | 2 | 3 | 4 | 5 | 6 | 7 |
| **All species (n = 10,999)** |  |  |  |  |  |  |  |  |
| Observed | 6084 | 2290 | 1062 | 667 | 406 | 252 | 153 | 85 |
| Expected | 4190 | 3987 | 1940 | 683 | 176 | 18 | 5 | 0 |
| **Songbirds (n = 6,603)** |  |  |  |  |  |  |  |  |
| Observed | 4367 | 1313 | 511 | 228 | 123 | 46 | 11 | 4 |
| Expected | 3497 | 2455 | 584 | 65 | 2 | 0 | 0 | 0 |
| **Non-songbirds (n = 4,396)** |  |  |  |  |  |  |  |  |
| Observed | 1717 | 977 | 551 | 439 | 283 | 206 | 142 | 81 |
| Expected | 693 | 1532 | 1356 | 618 | 174 | 18 | 5 | 0 |
| **All species, Nearctic + Neotropical (n = 4,648)** |  |  |  |  |  |  |  |  |
| Observed | 2737 | 782 | 398 | 315 | 198 | 112 | 63 | 43 |
| Expected | 1731 | 1683 | 842 | 299 | 82 | 9 | 2 | 0 |
| **All species, Afrotropical (n = 2,372)** |  |  |  |  |  |  |  |  |
| Observed | 1260 | 493 | 229 | 156 | 99 | 64 | 45 | 26 |
| Expected | 889 | 884 | 418 | 141 | 32 | 8 | 0 | 0 |
| **All species, Palearctic (n = 1,874)** |  |  |  |  |  |  |  |  |
| Observed | 656 | 493 | 292 | 180 | 110 | 69 | 44 | 30 |
| Expected | 696 | 651 | 360 | 131 | 34 | 2 | 0 | 0 |
| **All species, S&SE Asia (n = 3,143)** |  |  |  |  |  |  |  |  |
| Observed | 1477 | 799 | 376 | 211 | 110 | 78 | 59 | 33 |
| Expected | 1193 | 1123 | 562 | 214 | 49 | 2 | 0 | 0 |
| **All species, Australasia + Oceania (n = 1,676)** |  |  |  |  |  |  |  |  |
| Observed | 929 | 370 | 163 | 90 | 53 | 26 | 26 | 19 |
| Expected | 572 | 593 | 337 | 139 | 29 | 3 | 3 | 0 |

The multi-way comparison between datasets, which compared observed and expected frequencies of congruence between lists for all 120 possible multi-way intersections between the seven datasets, yielded similar results; in every case, the number of species that were shared between lists was significantly higher than would be expected by chance (in all cases, *p* < 10^-10^; Supplementary Figure S1).

For each of the six datasets that yielded quantitative measures of trade volume, there was a strong positive association between the frequency of recording (number of incidents) of species within each dataset and the frequency with which they were recorded in the remaining six datasets (Figure 2). A similar but generally weaker association was apparent for the total numbers of individual birds across the five datasets for which count data were available (Supplementary Figure S2). Thus there was strong evidence that the frequency with which species were recorded across datasets captures information on their relative abundance within datasets.

**Figure 2.**
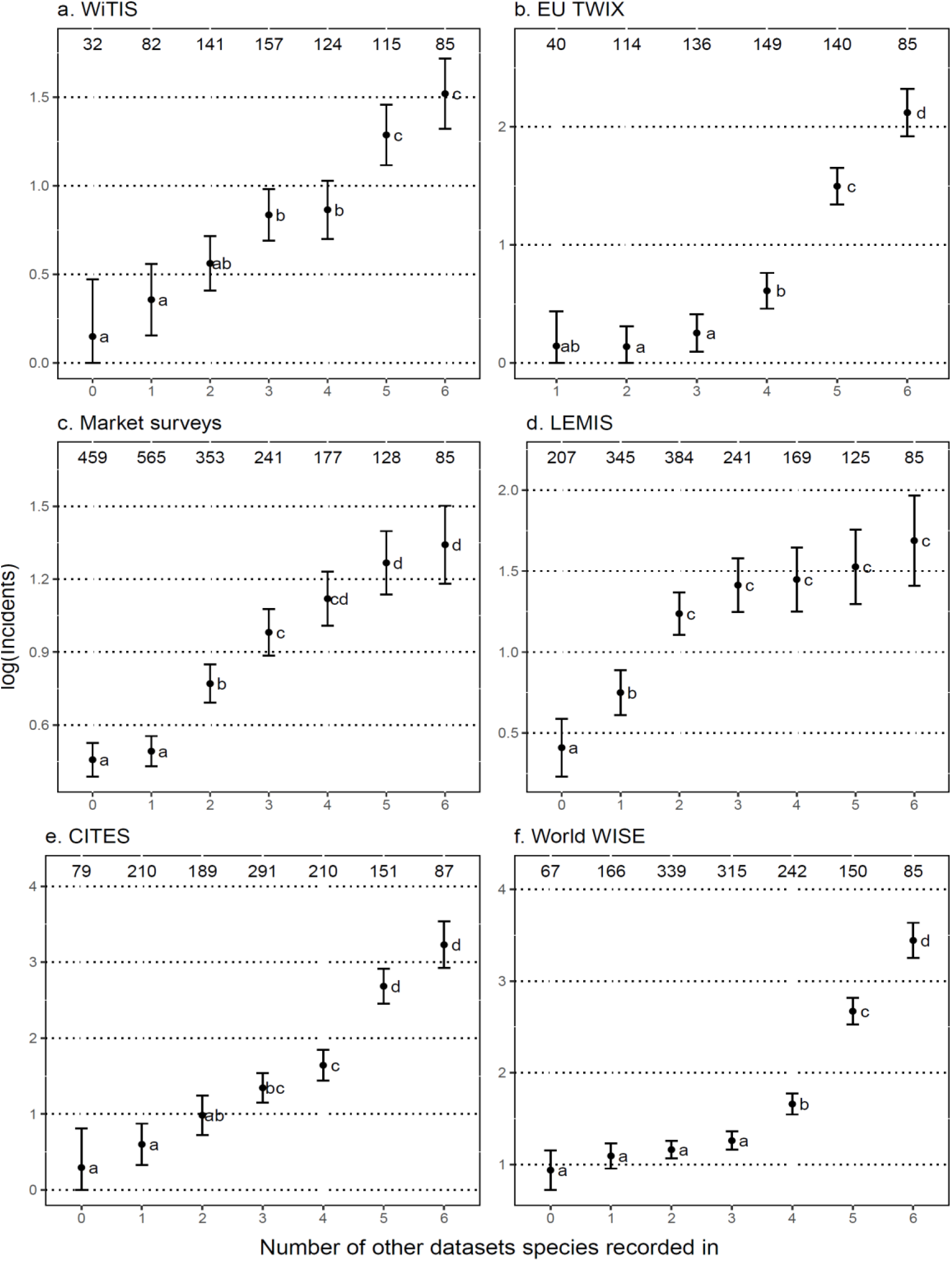
Relationship between frequency of reporting (mean log number of incidents, transactions or reports +/− 1 SE) of species within each of six quantitative trade databases and the number of other trade datasets that species were recorded in (range 0 to 6, including the Red List dataset). Numbers above the bars indicate the number of species in each case. In the case of EU TWIX, only three species were recorded in no other trade datasets, so these were merged with species occurring in one other trade dataset. Groups of species sharing a common letter did not differ at *p* < 0.05 (ANOVA with *post hoc* Tukey tests). The abbreviated names of the datasets follow those shown in Table 1.

The 265 species considered by independent studies or expert opinion to be heavily or unsustainably traded had a median trade prevalence score of 3, and 256 species (96.6%) were recorded in at least one of the seven global datasets, compared to a median score of 0 across all other species and a median of 2 across all other species that were recorded in trade at least once (Wilcoxon rank tests, *p* < 10^-12^ in both cases; Figure 3a). All of the 50 species whose trade has been the subject of published papers or reports were recorded at least once in the seven trade datasets (median = 3 datasets, range = 1-7; Supplementary Table S1). Similarly, the 202 species identified by Challender *et al*. (2023) as being threatened by international trade had significantly higher trade prevalence scores (median = 3) than all other species (median = 0; Wilcoxon rank test, *p* < 10^-12^), and all 27 priority species of the Asian Songbird Trade Specialist Group (2022) were recorded at least once (median = 3 compared to a median of 0 for all other species; Wilcoxon rank test, *p* < 10^-12^).

**Figure 3.**
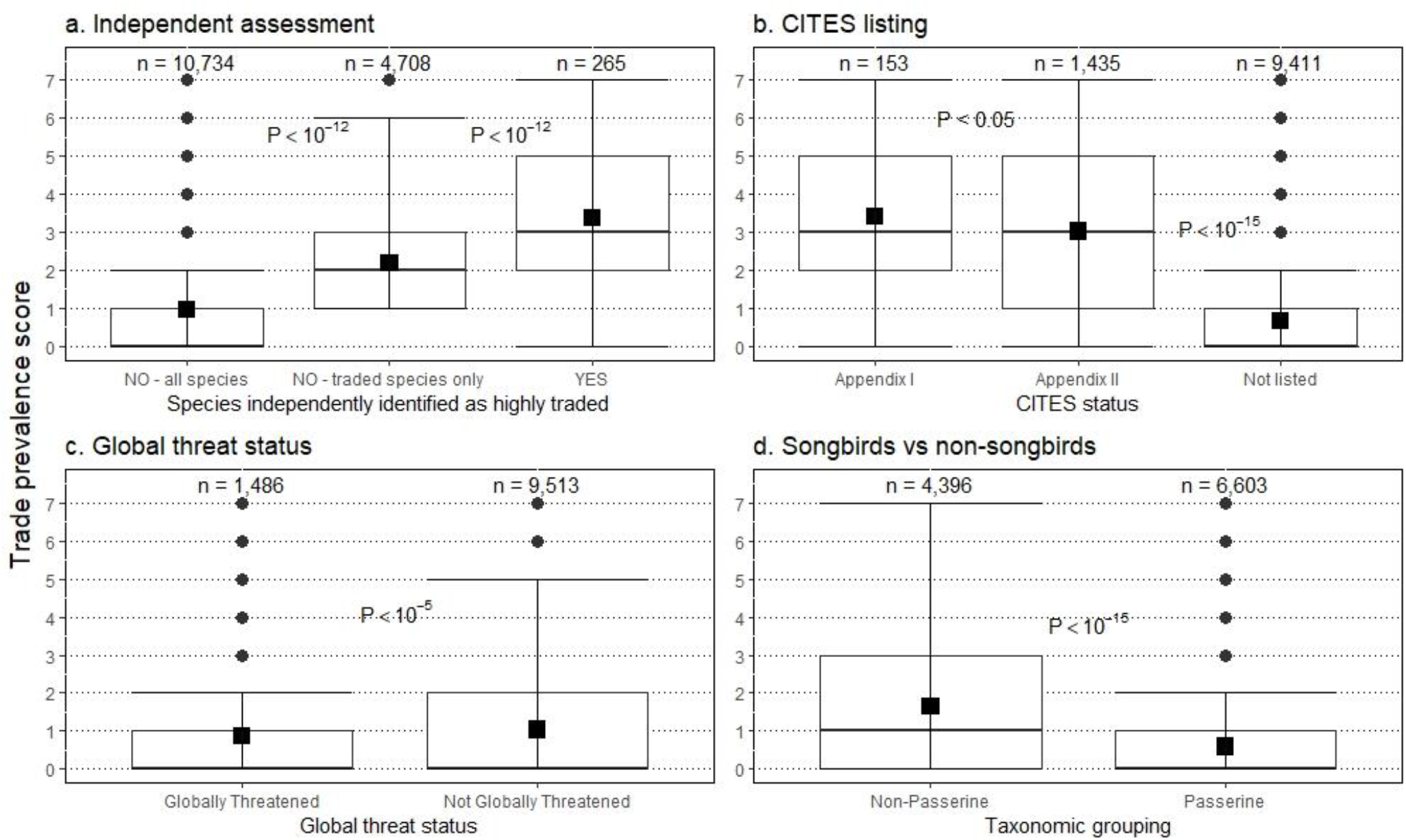
Relationship between the 7-level trade prevalence score (the number of datasets a species was recorded in) and (a) whether or not the species was independently identified as being heavily and perhaps unsustainably traded (comparison with all other species, and only those other species recorded in at least one dataset), (b) CITES Appendix listing (with the two largely CITES focused datasets removed, so maximum score = 5), (c) global threat status and (d) broad taxonomic division (songbirds *vs* all other orders). P-values show the results of Wilcoxon tests of the groupings on either side. In 3b, the P-value indicating the difference between CITES Appendix II-listed species and species not listed in CITES appendices was based on a comparison of trade prevalence scores that excluded the two datasets that contain data largely on CITES-listed species (i.e. it was based on a comparison of scores that run from 0 to 5).

We therefore conclude that the total number of datasets in which a species was recorded (0 to 7) is a broadly representative score of its relative prevalence in trade (with caveats discussed further below), although not necessarily of the extent to which it is threatened by trade.

### Prevalence in trade and its correlates

Univariate tests identified a number of significant predictors of the trade prevalence score. Species listed in CITES Appendices I or II had significantly higher trade prevalence scores on average than species not listed, and species listed in Appendix I had significantly higher scores than those listed in Appendix II (Figure 3b). The two trade datasets that comprised largely CITES-listed species (CITES and World WISE) were excluded from the trade prevalence score when statistically comparing CITES *vs* non-CITES species.

There was a strongly positive relationship between the trade prevalence score and extent of occurrence (ANOVA: F_7, 10,991_ = 152.5, *p* < 10^-16^). Species not listed as globally threatened had significantly higher median trade prevalence scores than those listed as globally threatened (i.e. Red List categories Vulnerable, Endangered, Critically Endangered; Figure 3c), although this may be a reflection of the fact that non-threatened species have significantly larger mean extent of occurrence than globally threatened species (t-test, *p* < 10^-16^).

The ordinal logistic regression model identified CITES status (t = 37.23, *p* < 10^-20^), broad taxonomic grouping (songbird vs non-songbird; t = 7.52, *p* < 10^-10^) and extent of occurrence (t = 32.69, *p* < 10^-15^), but not generation length (t = −0.55, *p* > 0.5), as significant predictors of trade prevalence. Species more likely to be recorded in higher trade prevalence classes tended to be CITES-listed non-songbirds with large geographical distributions (Figure 4).

**Figure 4.**
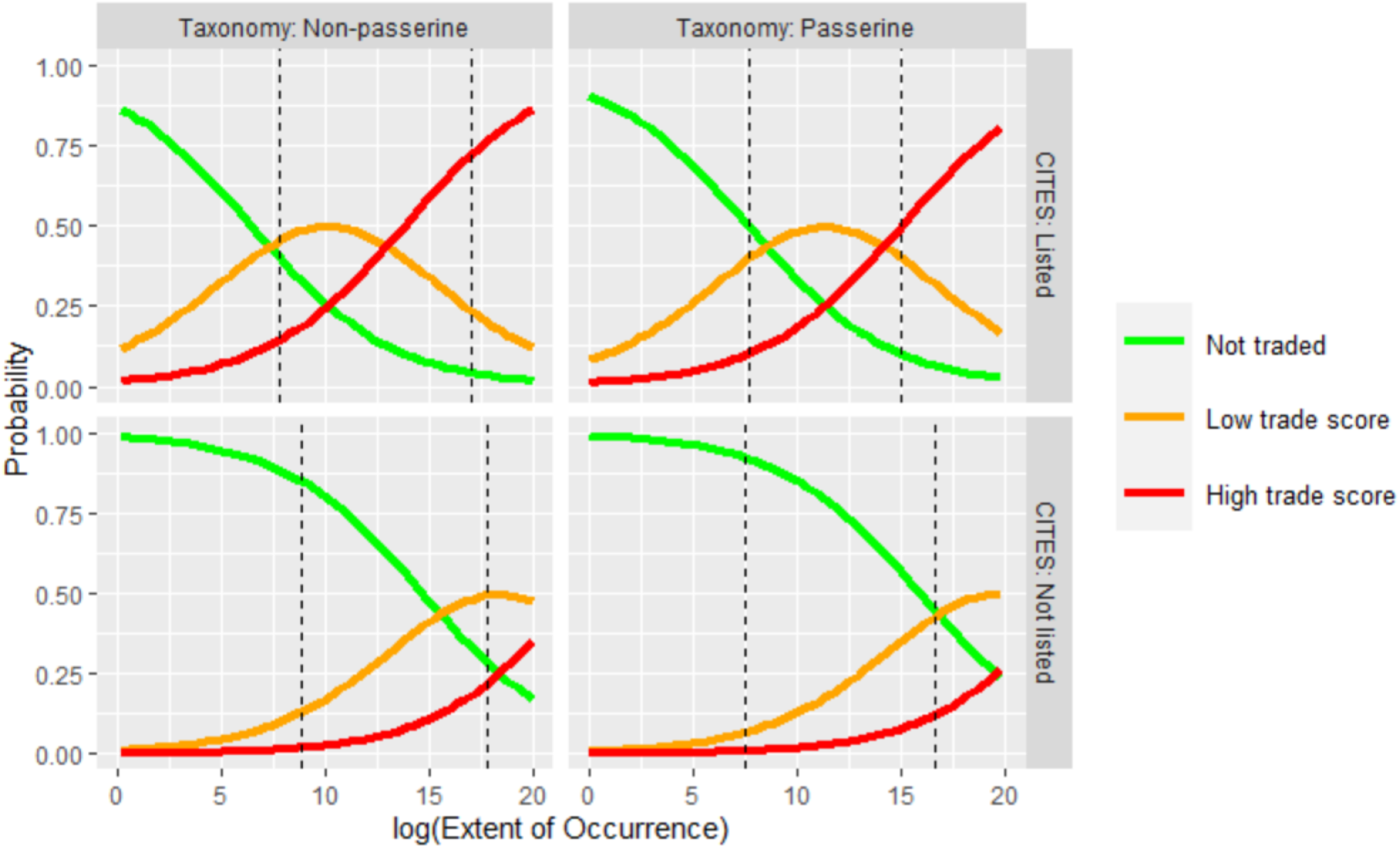
Predicted values, back-transformed to probabilities, from an ordinal logistic regression model of an ordered three-level trade prevalence factor with three predictors: broad taxonomic grouping (songbirds *vs* other orders), CITES status (listed in Appendix I or II, or not listed) and geographical distribution (extent of occurrence, EOO). The vertical dashed lines indicate the 5^th^ and 95^th^ percentiles of EOO for each subset of species (i.e. 90% of all species in each panel fall between the two vertical lines).

Songbirds (order Passeriformes) had a significantly lower median trade prevalence score than all other avian orders combined (Figure 3d) and the lowest among any speciose order except the Procellariiformes (albatrosses, petrels and shearwaters) (Table 2). However, songbirds comprise 60% of all bird species and the low mean prevalence score alone presents a misleading picture in terms of the number of species in trade. Songbirds comprised 45% of all species that were recorded in trade at least once, and were second only to parrots in terms of the number of species recorded in at least four of the trade datasets (Table 2). Of the 184 songbird species with trade prevalence scores of 4 or above, only 25 (13.6%) are listed in CITES Appendices I or II (and only two, Bali myna *Leucopsar rothschildi* and red siskin *Spinus cucullatus*, are listed in Appendix I), compared with 83.8% of the 794 non-songbird species with equivalent scores. Of the 133 globally threatened songbirds recorded in trade at least once, only 21 (15.8%) are listed in CITES Appendices I and II, compared to 277 (71.9%) of 393 globally threatened non-songbird species (Figure 5).

**Figure 5.**
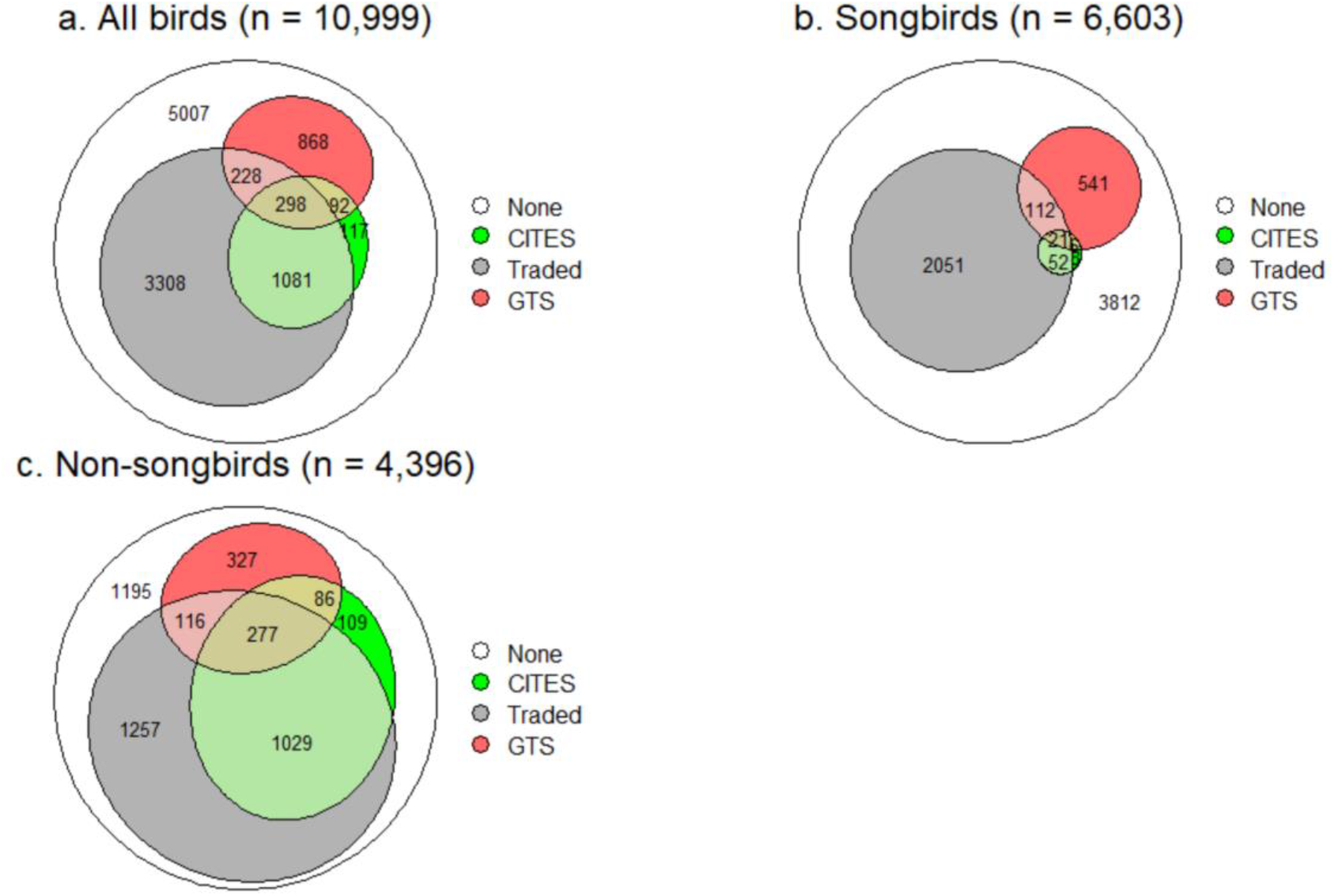
Area-proportional Venn diagrams showing the intersection between traded species (those recorded in at least one dataset) (‘Traded’), species listed in CITES Appendices I and II (‘CITES’) and globally threatened species (‘GTS’) for (a) all birds and for (b) songbirds and (c) non-songbirds separately. The white portion of each diagram (‘None’) represents species that were not recorded in trade, not listed in the CITES Appendices and not globally threatened.

### Geographical distribution of traded species

The geographical distributions of different groups of species with high trade indices are shown in Figure 6. These maps show hotspots of species prevalent in trade, not necessarily where trade pressure is highest. High numbers of species with trade prevalence scores of 5 or above occur in southern Brazil and eastern Paraguay, across Africa just south of the Sahara and through eastern Africa, and in southern Asia (Figure 6a). When only songbirds with high trade prevalence scores were considered, hotspots of co-occurring species were apparent in South-East Asia and in a broad band across the Palearctic (Figure 6b). Because most of the species with the highest trade prevalence scores were non-songbirds, a map of hotspots of non-songbirds (Figure 6c) was very similar to that for all species with the highest trade prevalence scores (Figure 6a). Similarly, because most CITES-listed species are non-songbirds, hotspots of CITES-listed species with high trade prevalence scores (Figure 6d) closely matched those of non-songbirds (Figure 6c), whereas hotspots for non-CITES species (Figure 6e) matched those of songbirds (Figure 6b).

**Figure 6.**
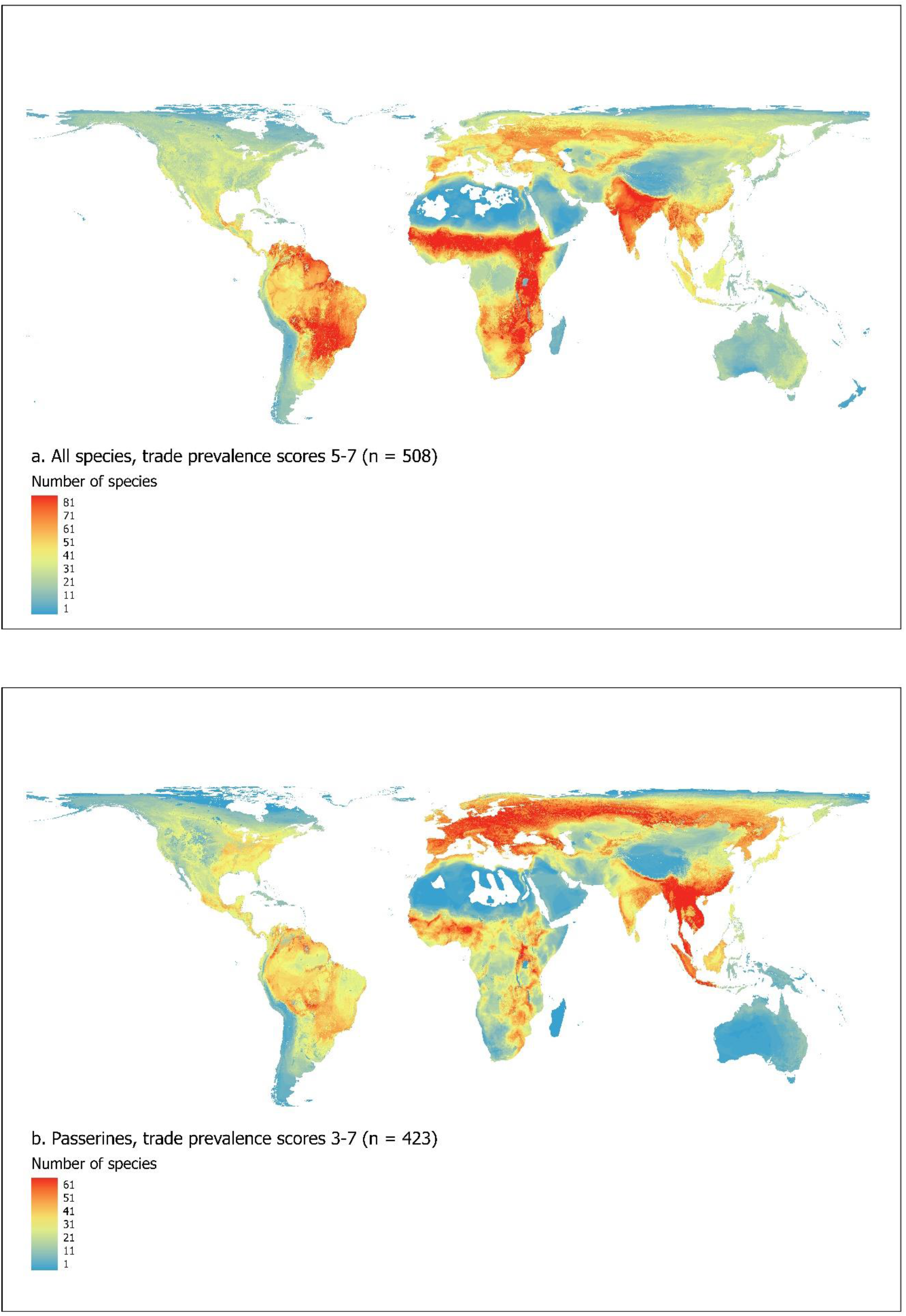
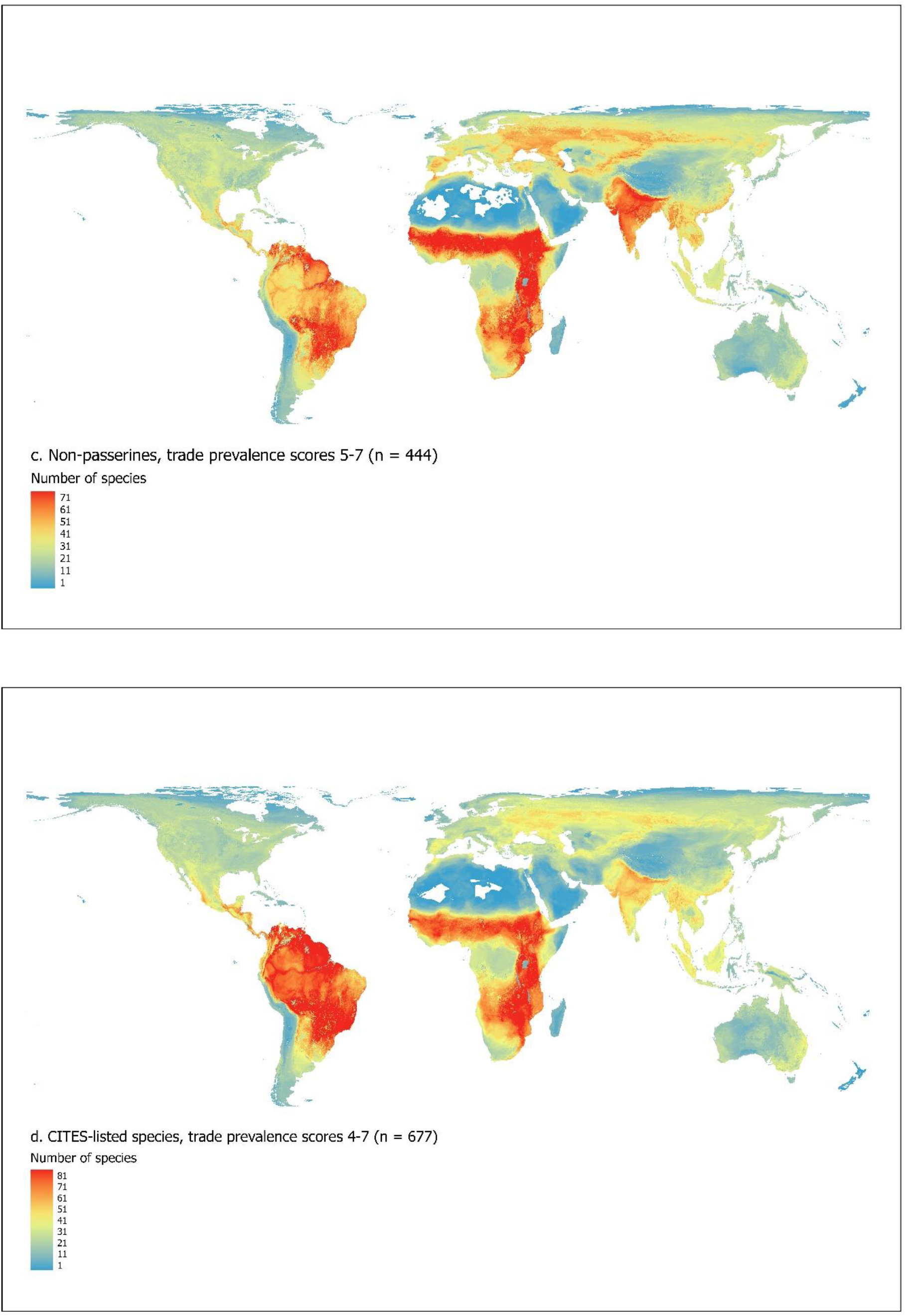
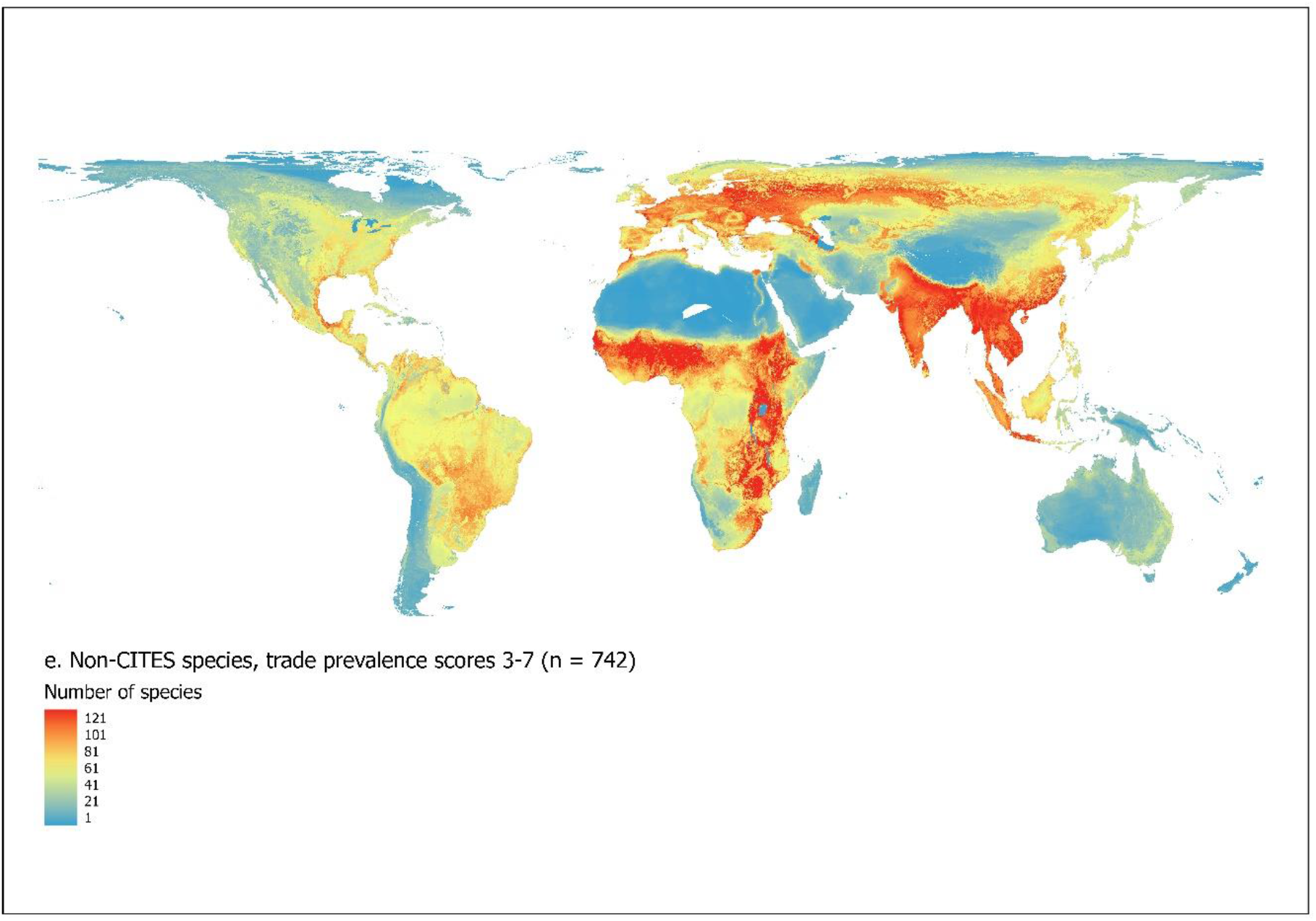
Maps showing number of (a) all species with trade prevalence scores of 5 or above, (b) songbirds with trade prevalence scores of 3 or above, (c) non-songbirds with trade prevalence scores of 5 or above, (d) CITES–listed species with trade prevalence scores of 4 or above and (e) non-CITES–listed species with trade prevalence scores of 3 or above. Thresholds vary between groups to equalise sample sizes. Sample sizes in each case are given in the figure legends. Colours indicate the numbers of traded species in each 5-km square, not necessarily hotspots of trade activity or volume of trade.

## DISCUSSION

Despite the very different methodologies and geographical and taxonomic foci of the seven trade datasets we examined, we found that species recorded in trade in one dataset were far more likely to be recorded in trade in one or more other datasets than would be expected by chance. The converse was, axiomatically, also true; species not recorded in one dataset were much less likely to be recorded in others than would be expected by chance. This pattern was strong at both global and regional scales, and for songbirds and non-songbirds separately. Furthermore, the frequency and numbers in which a species was recorded within datasets was positively correlated with its occurrence in other datasets, and species identified independently as being heavily and/or unsustainably traded were recorded significantly more frequently across datasets than the average across all other birds. This gives us confidence that heavily traded species were likely to have been recorded at least once across the seven datasets, and that the number of datasets a species was recorded in, which we term the trade prevalence score, can be used as a broad indicator of its prevalence in trade relative to other species. Furthermore, the high level of congruence between the global trade datasets, which between them document what is likely to be only a small proportion of the overall volume of trade actually taking place, suggests that they generate a consistent picture of the underlying patterns. Of the seven datasets we used, three record only illegal trade, two record only legal trade and two cover both illegal and legal trade, yet there was strong congruence between all of them in the species recorded. Our results therefore support previous suggestions that legal and illegal trade are broadly correlated (Olsen *et al*. 2019, Tittensor *et al*. 2020).

There are a number of caveats attached to the trade prevalence scores, particularly at the level of individual species. It is entirely possible that some sectors of the global trade in wild birds are under-represented or even missing altogether from the datasets we used, and therefore that the trade prevalence scores of some species are under-estimated. In particular, local trade in birds for medicine, ornaments or food may be greatly underestimated in comparison to longer-distance trade in birds as pets. Of the 265 species independently identified as being heavily or unsustainably traded, nine (3.4%) were not recorded in any of the seven datasets (Table S2). Similarly, five (2.5%) of the 202 species identified by Challender *et al*. (2023) as likely to be unsustainably traded were not recorded in any dataset. In some cases, these were likely to be due to taxonomic mismatches, as they relate to species that have been recognised only recently or are not recognised by all taxonomies. For example, the five unsustainably traded species listed by Challender *et al*. that have a trade prevalence score of zero in our analyses include *Amazona guatemalae*, *Eclectus cornelia* and *Polyplectron schleiermacheri*, which are either recently or not universally recognised. Their parent species, as which they would have been recorded in most of the datasets we used, all have high trade prevalence scores (respectively, *Amazona farinosa* [7], *Eclectus roratus* [6] and *Polyplectron malacense* [3]).

Conversely, some species might have scores that inflate the prevalence of wild-caught individuals in trade because, while we excluded captive-bred birds where possible, it was not possible to exclude them entirely; for example, budgerigar *Melopsittacus undulatus*, a widely captive-bred parrot that is explicitly excluded from CITES Appendices, attained a score of 5. Furthermore, all trade datasets are potentially flawed by the incorrect identification of species and by different and inconsistent (within and between datasets) taxonomic concepts. Incorrect identification is likely to affect some groups more than others; for example white-eyes (Zosteropidae) are much in demand in some trade sectors but identifying individuals to species is often problematic and birds may be entered in the data at higher taxonomic levels, meaning that the trade prevalence scores for individual species may be underestimated. We aligned all the datasets to a single global taxonomic list of birds, but our translation of taxonomic concepts is likely to be imperfect. Some datasets were gathered from multiple sources over a wide span of years using unstandardized taxonomies, so it is not always clear what taxonomic concept was being reported, and some concepts within a single dataset may have overlapped or changed over time. However, the use of a simple presence/absence metric summed across datasets might reduce some of the problems that have been identified in using quantitative data from large global trade datasets, such as the different time periods over which species have been recorded, the use of different and non-comparable recording units (whole birds, body parts, feathers etc.) and the multiple reporting within datasets of the same individuals (Robinson & Sinovas 2018).

Our results indicate that 45% of all bird species are recorded in trade, and that species listed in CITES Appendices I and II are significantly more prevalent in trade than non-CITES species. This suggests that CITES Appendices are successful at capturing many of the bird species in highest demand in trade. However, because CITES-listed species are afforded greater visibility and legal grounds for enforcement effort, and therefore higher detection in trade, they may be over-represented even in datasets whose focus is not explicitly CITES species. Nevertheless, our analyses also identified taxonomic groups that have high mean trade prevalence scores but low representation in CITES Appendices, such as the Anseriformes, Pelecaniformes, Galliformes and Columbiformes (Table 2). Furthermore, while only 1.3% songbird species are listed in CITES Appendices, songbirds accounted for a 45.5% of all traded species and 20.5% of species with trade prevalence scores of 4 or higher (Table 2). Although 86.8% of CITES-listed species were recorded at least once in trade, they accounted for only 28.1% of all traded species (Figure 5), suggesting that our results are not unduly influenced by the relative over-reporting of CITES species.

Of the 896 species with the highest trade prevalence scores (4 or above), representing around 8% of the world’s species, 298 (33.3%) are not listed in CITES Appendices I or II, and 11 are globally threatened. These are great curassow *Crax rubra* (Vulnerable), bare-faced curassow *Crax fasciolata* (Vulnerable), highland guan *Penelopina nigra* (Vulnerable), helmeted curassow *Pauxi pauxi* (Endangered), lesser adjutant *Leptoptilos javanicus* (Vulnerable), southern ground-hornbill *Bucorvus leadbeateri* (Vulnerable), European turtle-dove *Streptopelia turtur* (Vulnerable), Javan leafbird *Chloropsis cochinchinensis* (Endangered), greater green leafbird *Chloropsis sonnerati* (Endangered), yellow-breasted bunting *Emberiza aureola* (Critically Endangered) and great-billed seed-finch *Sporophila maximiliani* (Endangered).

Mapping the trade prevalence scores suggests that large numbers of heavily traded non-CITES species co-occur in sub-Saharan Africa, South and South-East Asia and across Eurasia (Figure 5e). Our results support recent concern about the trade in songbirds in South-East Asia, confirming this region as a hotspot of songbird species with high trade prevalence scores (Figure 5b). However, we also identify Eurasia and parts of Africa as regions holding large numbers of songbirds with high trade prevalence scores. Nearly half (45.5%) of all the species recorded in trade, and a fifth (20.9%) of those with scores of 4 or above, were songbirds, yet only 1.3% of all songbirds are listed in CITES Appendices I and II (Table 2).

To determine whether a species should be included in CITES Appendices I or II, biological and trade criteria are applied (CITES 2016). Appendix I criteria concern extinction risk brought about by small populations or restricted areas of distribution, in either case exacerbated by additional vulnerability factors such as numbers of occupied sites, fluctuations and marked declines. Appendix II is appropriate when regulation of trade in a species is necessary to avoid its status deteriorating, to prevent it becoming eligible for inclusion in Appendix I or to ensure that the wild harvest remains sustainable. There are no such criteria for Appendix III listing, reflecting its rather different function. Prevalence in trade *per se* is not part of any criterion, but our scores could be used as a filter to prioritise the identification of species to be considered for future CITES listing or, where already listed, for transfer between Appendices. CITES Parties review which species should be newly listed, or transferred, as with recent concerns about the inadequate representation of songbirds in the Appendices (supported by our analysis). However, identifying the set of species finally to be listed is challenging. Of the 490 bird species with the highest trade prevalence scores (5, 6 or 7), 98 (20.0%) are not listed in CITES Appendices I or II (Table S4), nearly half of them songbirds. These could form a short-list of taxa for subsequent analysis applying the CITES listing criteria, to be integrated with other outputs such as the priority taxa list of the Asian Songbird Trade Specialist Group (2022). There probably can never be a simple, all-encompassing route to identifying which species to list, and each taxon needs to be assessed and reviewed individually. Some taxa in need of protection under CITES may have low prevalence scores, while some with high scores may not benefit from or merit CITES listing, either because trade is of little conservation concern or because it is predominantly domestic, and thus falls outside of the focus of the convention.

This study highlights the substantial extent of the trade in wild birds, which affects almost half of all species, and suggests there is a need to expand CITES listing, in particular for songbirds and some other widely traded orders of birds. Urgent action is needed to monitor and address unsustainable and illegal trade using a variety of tools from enforcement of legislation to promotion of alternative livelihoods and demand reduction. International intergovernmental task forces have been established to provide guidance, support and experience sharing, such as the CMS Intergovernmental Task Force on Illegal Killing, Taking and Trade of Migratory Birds in the Mediterranean (MIKT) and the Intergovernmental Task Force to Address Illegal Hunting, Taking and Trade of Migratory Birds in the East Asian-Australasian Flyway (ITTEA). Signatories of the Bern Convention and CMS MIKT are working towards achieving the goals of the Rome Strategic Plan and countries in the Middle East are working together to achieve the goals of a Roadmap to tackle the issues around illegal and unsustainable take and trade. Insights from criminology and social science can be used to reduce illegal trade and to support a better distribution of potential benefits from sustainable use or alternative livelihoods (Pires *et al*. 2016, Pires *et al*. 2021)

Our methodology for developing a broad metric of trade that can be applied across an entire taxonomic class could be expanded for birds, and extended to other taxonomic groups, by incorporating new sources of data into the calculation of the score, for example through data gathered from online markets and social media (Di Minin *et al*. 2019, Davies *et al*. 2022, Okarda *et al*. 2022). This could be used to help prioritise species for Red List assessments or updates. However, it is important to recognise that the trade prevalence score is not a metric of the extent to which species are threatened by trade. New approaches are required if we are to assess in a systematic way the extent to which trade threatens the survival of traded species.

## ACKNOWLEDGEMENTS

This work was supported by the CCI Collaborative Fund, which is funded by Arcadia (a charitable fund of Lisbet Rausing and Peter Baldwin), the Rothschild Foundation, the A.G. Leventis Foundation, the Isaac Newton Trust and the Prince Albert II of Monaco Foundation. We thank the many thousands of individuals and organisations who contribute to BirdLife’s assessments of extinction risk for all the world’s birds for the IUCN Red List. For helpful comments and the provision of additional data we thank Alex Berryman, Nigel Collar, Jacqueline Jürgens, LoraKim Joyner (One Earth Conservation), Boleslaw Slocinski (Association Biom), Jihad, Ferry Hasudungan, Achmad Ridha Junaid (Burung Indonesia), Stella Egbe (Nigerian Conservation Foundation), Guy Shorrock (RSPB), Jarryd Alexander and members of the Asian Songbird Trade Specialist Group (ASTSG): David Jeggo, Sofiya Shukhova, Jessica Lee, Andrew Owen, Stuart Marsden, Frank Rheindt, Simon Bruslund, Bas Van Balen, Sicily Fiennes, Laura Gardner, Rosa Gleave and James Eaton.

