## Supplementary data for "Assessing the global prevalence of wild birds in trade"

### **Supplementary Data:** "Assessing the global prevalence of wild birds in trade"

**List of Contents**

**Figure S1.** (p. 2) Observed frequency with which species were recorded in (S1a) two, (S1b) three, (S1c) four or (S1d) five to seven trade datasets.

**Figure S2.** (p. 5) Relationship between numbers of individual birds (log number of individuals +/- 1 SE) of species within each of five quantitative trade databases and the number of other trade databases that species were recorded in.

**Table S1**. (p. 6) List of 50 species known from sources other than the seven global datasets to be heavily traded, and the number of those datasets each species was recorded in.

**Table S2**. (p. 14) List of species reported as being heavily or unsustainably traded in questionnaire surveys and/or publications and the number of datasets each was recorded in (trade prevalence score).

**Table S3.** (p. 23) List of publications used to compile the dataset on birds in markets and other sources.

**Table S4.** (p. 29) The 98 bird species with trade prevalence scores of 5 or above that are not listed in CITES Appendices I or II.

**Figure S1**

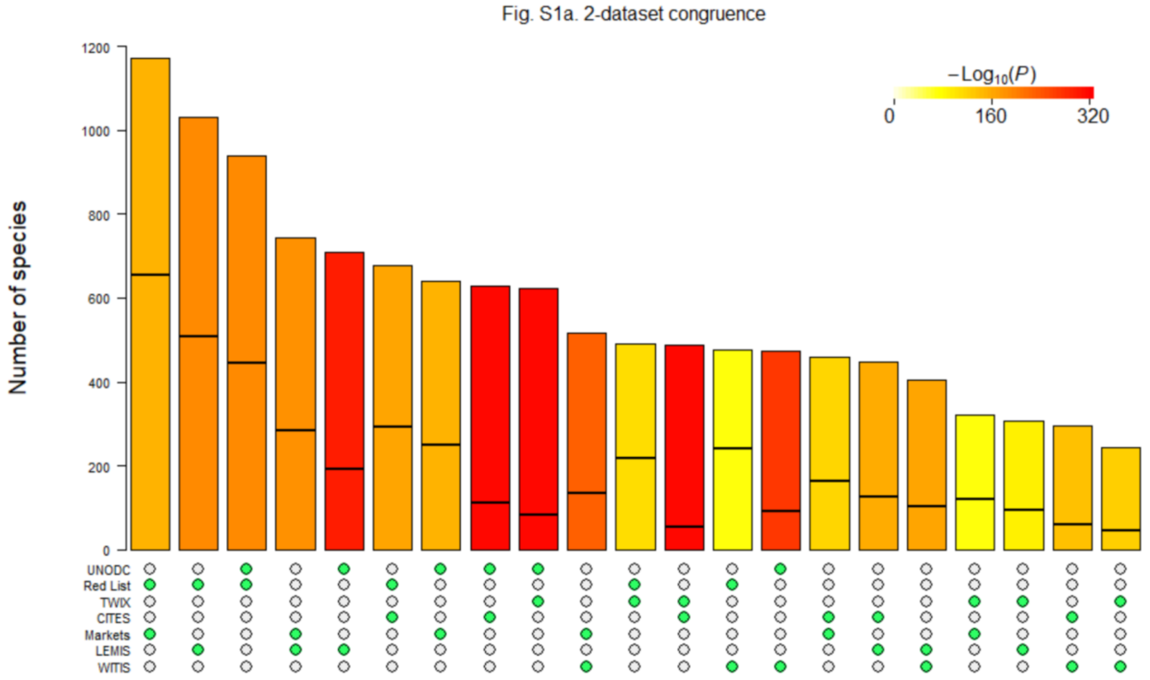

Fig. S1a. 2-dataset congruence
−Log10(P)
0
160
320
Number of species
1200
1000
800
600
400
200
0
UNODC
Red List
TWIX
CITES
Markets
LEMIS
WITIS

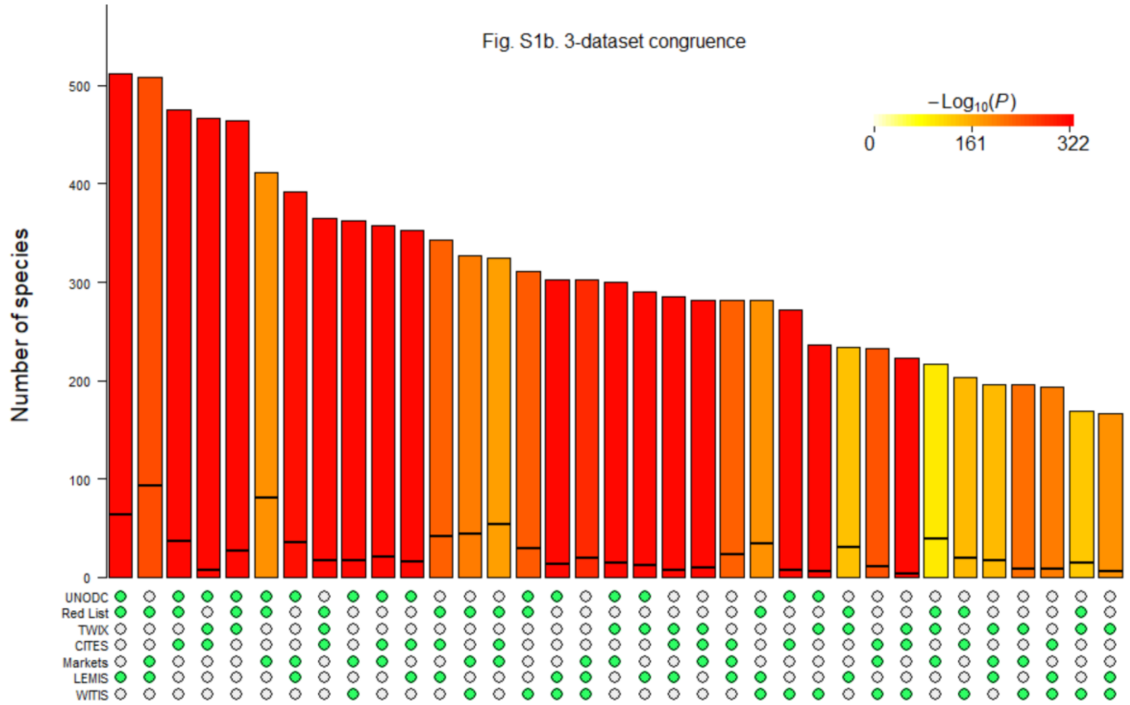
Fig. S1b. 3-dataset congruence
−Log10(P)
0
161
322
Number of species
0
100
200
300
400
500
UNODC
Red List
TWIX
CITES
Markets
LEMIS
WITIS

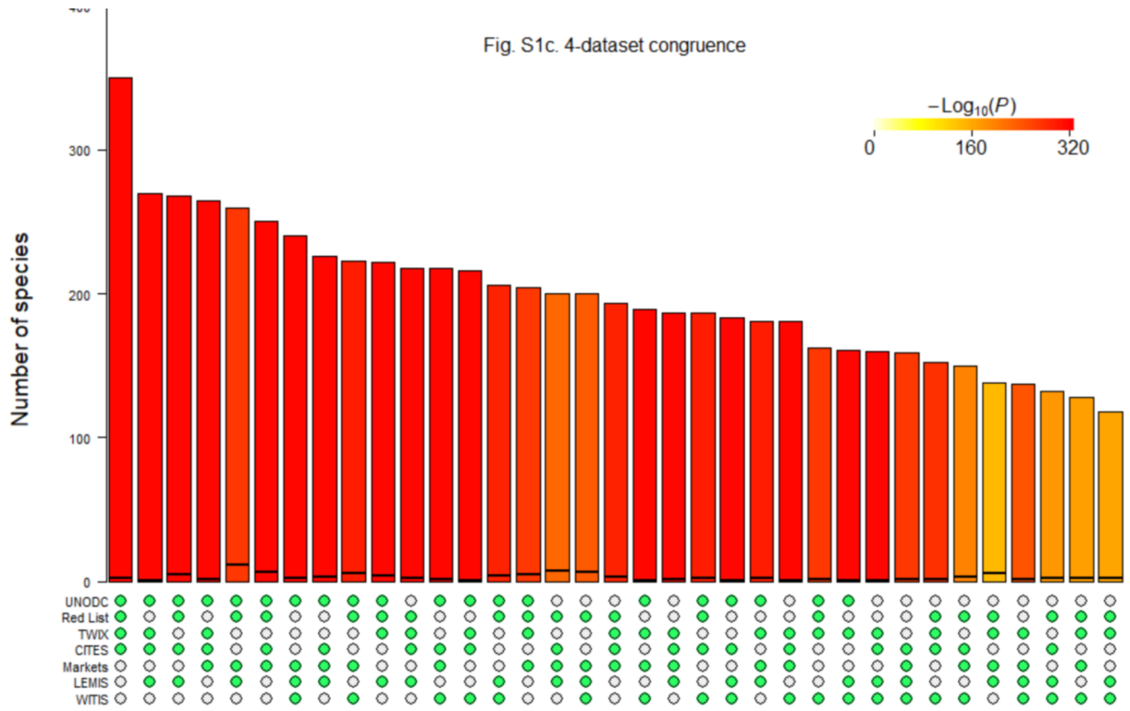
Fig. S1c. 4-dataset congruence
−Log10(P)
0
160
320
Number of species
0
100
200
300
UNODC
Red List
TWIX
CITES
Markets
LEMIS
WITIS

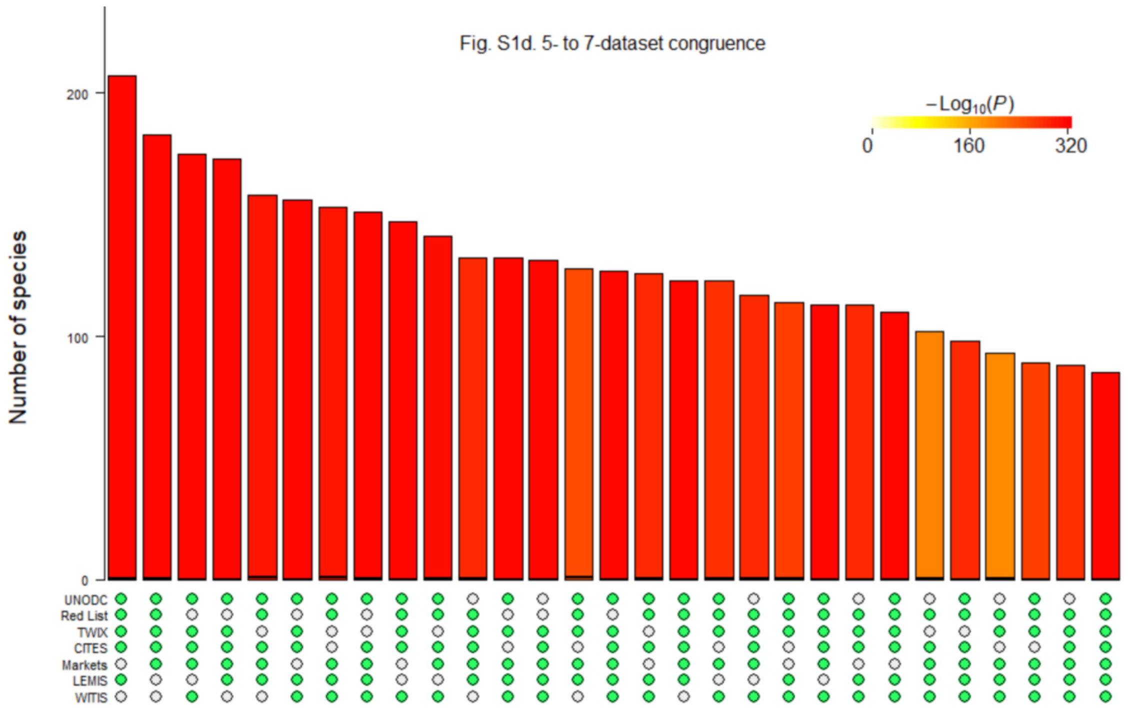

**Figure S1.** Observed frequency with which species were recorded in (S1a) two, (S1b) three, (S1c) four or (S1d) five to seven trade datasets. The expected frequency under the null hypothesis of no association, based upon the number of species recorded in each dataset and the number of bird species globally, is shown as a horizontal black line (which lies close to zero in intersections between four or more datasets). Bar colour indicates the probability of the observed frequency being obtained under the null hypothesis of no association; in all cases, $P < 10^{-14}$. Graphs produced using the R package *SuperExactTest* (Wang *et al*. 2015). Abbreviated dataset names are explained in Table 1 of the main paper.

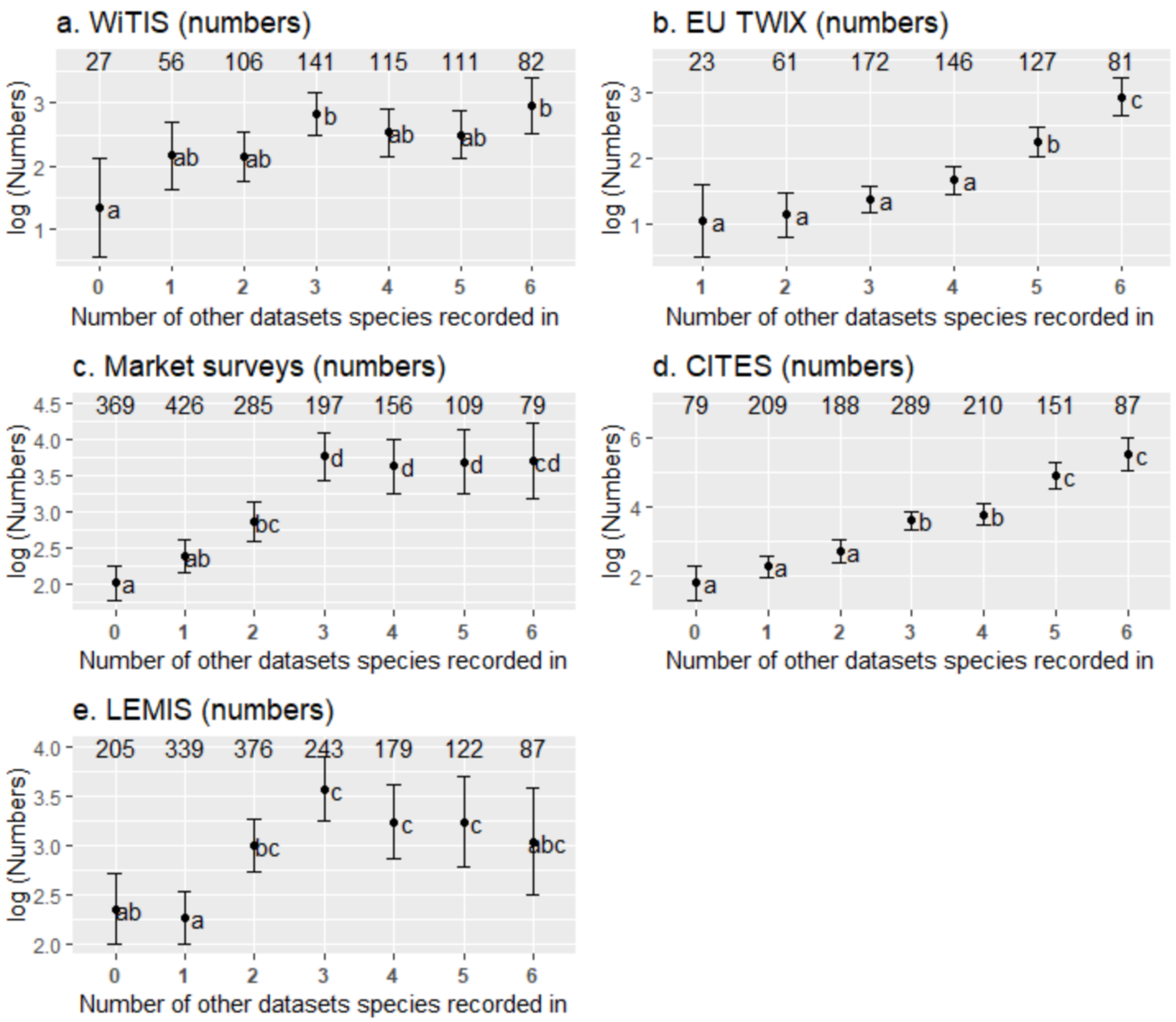

**Figure S2.** Relationship between numbers of individual birds (log number of individuals +/- 1 SE) of species within each of five quantitative trade databases and the number of other trade databases that species were recorded in (range 0 to 6, including the Red List dataset which does not contain data on frequency of occurrence). Numbers above the bars indicate the number of species in each case. In the case of EU TWIX, only three species were recorded in no other trade datasets, so these were merged with species occurring in one other trade dataset. Groups of species not sharing a common letter differed at $P < 0.05$ (ANOVA with *post hoc* Tukey tests). The abbreviated names of the datasets follow those shown in Table 1 of the main paper.

**Table S1.** List of 50 species assessed from sources other than the seven global datasets to be heavily or unsustainably traded, and the number of those datasets each species was recorded in. Those marked with an asterisk by the common name were reported in questionnaire surveys, those with the common name <u>underlined</u> were identified as potentially threatened by trade by Challender *et al.* (2023), those with the common name in *italics* (songbirds only) were identified as potentially threatened by trade by Juergens *et al.* (2021), those with the common name in **bold** (Asian songbirds only) are Tier 1 priority species of the IUCN Asian Songbird Trade Specialist Group (ASTAG). Species are listed by region in descending order of the number of global datasets they were recorded in.

| Scientific name | Common name | Region | No. datasets | References |
|---|---|---|---|---|
| *Necrosyrtes monachus* | Hooded Vulture | Afrotropics | 7 | Ogada & Buij (2011), Daboné *et al.* (2022) |
| *Psittacus erithacus* | <u>Grey Parrot</u>* | Afrotropics | 6 | Juste (1996), Ngenyi *et al.* (2003), Eniang *et al.* (2008), Hart *et al.* (2015), Marsden *et al.* (2015), Annorbah *et al.* (2016), Poole & Shepherd (2016), Martin (2018), Martin *et al.* (2018), Valle *et al.* (2018), Martin *et al.* (2019), Atoussi *et al.* (2020), Assou *et al.* (2021), Davies *et al.* (2022) |
| *Gyps coprotheres* | Cape Vulture* | Afrotropics | 5 | Beilis & Esterhuizen (2005) |
| *Balearica pavonina* | Black Crowned Crane* | Afrotropics | 5 | Kone *et al.* (2007) |
| *Psittacus timneh* | <u>Timneh Parrot</u>* | Afrotropics | 4 | Martin (2018), Valle *et al.* (2019), Atoussi *et al.* (2020) |
| *Poicephalus robustus* | Cape Parrot | Afrotropics | 3 | Pillay *et al.* (2010), Coetzer *et al.* (2017), Collar & Fishpool (2017) |
| *Vultur gryphus* | Andean Condor | Neotropics | 7 | Williams *et al.* (2012) |
| *Spinus cucullatus* | <u>*Red Siskin*</u> | Neotropics | 7 | Sánchez-Mercado *et al.* (2020) |
| *Eupsittula canicularis* | Orange-fronted Parakeet | Neotropics | 7 | Padilla-Jacobo *et al.* (2021) |
| *Brotogeris versicolurus* | White-winged Parakeet | Neotropics | 6 | Daut *et al.* (2016) |
| *Psittacara erythrogenys* | <u>Red-masked Parakeet</u>* | Neotropics | 6 | Best *et al.* (2010) |
| *Brotogeris pyrrhoptera* | <u>Grey-cheeked Parakeet</u>* | Neotropics | 2 | Best *et al.* (2010) |
| *Carduelis carduelis* | European Goldfinch* | Palearctic | 4 | Louadj *et al.* (2021) |

| | | | | |
|---|---|---|---|---|
| *Cacatua sulphurea* | Yellow-crested Cockatoo* | S&SE Asia | 7 | Cahill *et al.* (2006), Reuleaux *et al.* (2022), |
| *Cacatua goffiniana* | Tanimbar Corella* | S&SE Asia | 7 | Haryoko *et al.* (2021) |
| *Rhinoplax vigil* | Helmeted Hornbill | S&SE Asia | 6 | Collar (2015), Beastall *et al.* (2016), Krishnasamy *et al.* (2016) |
| *Cacatua alba* | White Cockatoo* | S&SE Asia | 6 | Lambert (1993) |
| *Lorius garrulus* | Chattering Lory* | S&SE Asia | 6 | Lambert (1993), Cottee-Jones *et al.* (2014) |
| *Garrulax canorus* | *Chinese Hwamei** | S&SE Asia | 6 | Shepherd *et al.* (2016a), Shepherd *et al.* (2020a), Nelson & Shepherd (2023) |
| *Gracula religiosa* | ***Common Hill Myna**** | S&SE Asia | 6 | Bernardo (2016), Ng *et al.* (2021) |
| *Copsychus saularis* | ***Oriental Magpie-robin**** | S&SE Asia | 5 | Chng *et al.* (2021) |
| *Garrulax leucolophus* | White-crested Laughingthrush | S&SE Asia | 5 | Shepherd (2007) |
| *Pycnonotus zeylanicus* | ***Straw-headed Bulbul*** | S&SE Asia | 5 | Shepherd *et al.* (2013), Chng *et al.* (2016), Bergin *et al.* (2017), Yong *et al.* (2017), Leupen & Shepherd (2018a), Chiok *et al.* (2019), Nijman *et al.* (2022) |
| *Leucopsar rothschildi* | ***Bali Myna**** | S&SE Asia | 5 | Jepson (2015), Nijman *et al.* (2017) |
| *Tanygnathus lucionensis* | Blue-naped Parrot | S&SE Asia | 4 | Bernardo (2016) |
| *Kittacincla malabarica* | ***White-rumped Shama**** | S&SE Asia | 4 | Ng *et al.* (2017), Leupen *et al.* (2018), Rheindt *et al.* (2019) |
| *Pycnonotus jocosus* | *Red-whiskered Bulbul* | S&SE Asia | 4 | Techachoochert & Round (2013) |
| *Chloropsis sonnerati* | ***Greater Green Leafbird**** | S&SE Asia | 4 | Chng *et al.* (2017) |
| *Pycnonotus goiavier* | *Yellow-vented Bulbul** | S&SE Asia | 4 | Nijman *et al.* (2022) |
| *Nisaetus bartelsi* | Javan Hawk-eagle* | S&SE Asia | 3 | Nijman *et al.* (2009) |
| *Pycnonotus bimaculatus* | *Orange-spotted Bulbul** | S&SE Asia | 3 | Leupen & Gomez (2019), Nijman *et al.* (2022) |
| *Acridotheres melanopterus* | ***Black-winged Myna**** | S&SE Asia | 3 | Shepherd *et al.* (2016b), Nijman *et al.* (2017), Nijman *et al.* (2018), Squires *et al.* (2022) |
| *Geokichla citrina* | ***Orange-headed Thrush**** | S&SE Asia | 3 | Kristianto & Jepson (2011) |
| *Arborophila javanica* | Chestnut-bellied Partridge | S&SE Asia | 2 | Nijman (2022) |
| *Garrulax bicolor* | ***Sumatran Laughingthrush**** | S&SE Asia | 2 | Shepherd (2007), Shepherd (2013), Shepherd *et al.* (2016a), Bušina *et al.* (2018), Bušina *et al.* (2020), Heinrich *et al.* (2021) |

| | | | | |
|---|---|---|---|---|
| *Garrulax mitratus* | *Chestnut-capped Laughingthrush** | S&SE Asia | 2 | Shepherd *et al.* (2016a) |
| *Acridotheres tricolor* | ***Grey-backed Myna**** | S&SE Asia | 2 | Bruslund *et al.* (2021) |
| *Arborophila orientalis* | White-faced Partridge* | S&SE Asia | 1 | Nijman (2022) |
| *Cissa thalassina* | ***Javan Green Magpie**** | S&SE Asia | 1 | Nijman *et al.* (2017) |
| *Garrulax rufifrons* | ***Rufous-fronted Laughingthrush**** | S&SE Asia | 1 | Collar & van Balen (2013), Shepherd *et al.* (2016a), Nijman *et al.* (2017), Nijman *et al.* (2020) |
| *Garrulax palliatus* | ***Sunda Laughingthrush**** | S&SE Asia | 1 | Shepherd *et al.* (2016a), Leupen *et al.* (2020) |
| *Laniellus albonotatus* | *Spotted Crocias** | S&SE Asia | 1 | Leupen & Shepherd (2018b), Nijman *et al.* (2021a) |
| *Gracupica jalla*[1] | ***Javan Pied Starling**** | S&SE Asia | 1 | van Balen & Collar (2021), Nijman *et al.* (2021b) |
| *Gracula venerata*[1] | ***Tenggara Hill Myna**** | S&SE Asia | 1 | Reuleaux *et al.* (2019) |
| *Otus jolandae*[1] | Rinjani Scops-owl* | S&SE Asia | 1 | Shepherd *et al.* (2020b) |
| *Falco cherrug* | Saker Falcon* | Multiple | 7 | Ma & Chen (2007), Dixon *et al.* (2011), Levin (2011), Shobrak (2015), Stretesky *et al.* (2018) |
| *Aquila fasciata* | Bonelli's Eagle | Multiple | 5 | Di Vittorio *et al.* (2018) |
| *Dendrocygna arcuata* | Wandering Whistling-duck* | Multiple | 4 | Nijman *et al.* (2022) |
| *Emberiza aureola* | *Yellow-breasted Bunting** | Multiple | 4 | Kamp *et al.* (2015) |
| *Turdus obscurus* | Eyebrowed Thrush | Multiple | 2 | Iqbal *et al.* (2018) |

[1]Species that have been taxonomically recognised only recently and therefore likely to have been recorded mostly under their parent species name in the global datasets.

**Table S2**. List of 265 species reported as being heavily or unsustainably traded in questionnaire surveys and/or publications and the number of datasets each was recorded in (trade prevalence score). Species listed in descending order of their trade prevalence score. Further details of the 50 species reported in independent studies ('Publication') are given in Table S1.

| Species | Common name | Source | Number of datasets (trade prevalence score) |
|---|---|---|---|
| *Accipiter gentilis* | Northern Goshawk | Questionnaire | 7 |
| *Amazona aestiva* | Turquoise-fronted Amazon | Questionnaire | 7 |
| *Amazona auropalliata* | Yellow-naped Amazon | Questionnaire | 7 |
| *Anodorhynchus hyacinthinus* | Hyacinth Macaw | Questionnaire | 7 |
| *Ara macao* | Scarlet Macaw | Questionnaire | 7 |
| *Aratinga solstitialis* | Sun Parakeet | Questionnaire | 7 |
| *Cacatua goffiniana* | Tanimbar Corella | Publication and Questionnaire | 7 |
| *Cacatua sulphurea* | Yellow-crested Cockatoo | Publication and Questionnaire | 7 |
| *Elanus caeruleus* | Black-winged Kite | Questionnaire | 7 |
| *Eupsittula canicularis* | Orange-fronted Parakeet | Publication | 7 |
| *Falco cherrug* | Saker Falcon | Publication and Questionnaire | 7 |
| *Falco peregrinus* | Peregrine Falcon | Questionnaire | 7 |
| *Haliaeetus leucogaster* | White-bellied Sea-eagle | Questionnaire | 7 |
| *Lonchura oryzivora* | Java Sparrow | Questionnaire | 7 |
| *Loriculus galgulus* | Blue-crowned Hanging-parrot | Questionnaire | 7 |
| *Lorius lory* | Black-capped Lory | Questionnaire | 7 |
| *Necrosyrtes monachus* | Hooded Vulture | Publication | 7 |
| *Pandion haliaetus* | Osprey | Questionnaire | 7 |
| *Probosciger aterrimus* | Palm Cockatoo | Questionnaire | 7 |
| *Psittacara leucophthalmus* | White-eyed Parakeet | Questionnaire | 7 |
| *Spinus cucullatus* | Red Siskin | Publication and Questionnaire | 7 |
| *Trichoglossus haematodus* | Coconut Lorikeet | Questionnaire | 7 |
| *Tyto alba* | Common Barn-owl | Questionnaire | 7 |
| *Vultur gryphus* | Andean Condor | Publication and Questionnaire | 7 |
| *Accipiter virgatus* | Besra | Questionnaire | 6 |

| | | | |
|---|---|---|---|
| *Amazona oratrix* | Yellow-headed Amazon | Questionnaire | 6 |
| *Ara chloropterus* | Red-and-green Macaw | Questionnaire | 6 |
| *Brotogeris versicolurus* | White-winged Parakeet | Publication | 6 |
| *Buteo buteo* | Eurasian Buzzard | Questionnaire | 6 |
| *Cacatua alba* | White Cockatoo | Publication and Questionnaire | 6 |
| *Cacatua galerita* | Sulphur-crested Cockatoo | Questionnaire | 6 |
| *Cacatua moluccensis* | Salmon-crested Cockatoo | Questionnaire | 6 |
| *Garrulax canorus* | Chinese Hwamei | Publication and Questionnaire | 6 |
| *Gracula religiosa* | Common Hill Myna | Publication and Questionnaire | 6 |
| *Haliastur indus* | Brahminy Kite | Questionnaire | 6 |
| *Lorius garrulus* | Chattering Lory | Publication and Questionnaire | 6 |
| *Milvus migrans* | Black Kite | Questionnaire | 6 |
| *Nisaetus cirrhatus* | Changeable Hawk-eagle | Questionnaire | 6 |
| *Paroaria coronata* | Red-crested Cardinal | Questionnaire | 6 |
| *Psittacara erythrogenys* | Red-masked Parakeet | Publication and Questionnaire | 6 |
| *Psittacus erithacus* | Grey Parrot | Publication and Questionnaire | 6 |
| *Rhinoplax vigil* | Helmeted Hornbill | Publication and Questionnaire | 6 |
| *Spilornis cheela* | Crested Serpent-eagle | Questionnaire | 6 |
| *Strix leptogrammica* | Brown Wood-owl | Questionnaire | 6 |
| *Alauda arvensis* | Eurasian Skylark | Questionnaire | 5 |
| *Amazona brasiliensis* | Red-tailed Amazon | Questionnaire | 5 |
| *Anas platyrhynchos* | Mallard | Questionnaire | 5 |
| *Aquila fasciata* | Bonelli's Eagle | Publication | 5 |
| *Ara ambiguus* | Great Green Macaw | Questionnaire | 5 |
| *Balearica pavonina* | Black Crowned Crane | Publication and Questionnaire | 5 |
| *Butastur indicus* | Grey-faced Buzzard | Questionnaire | 5 |
| *Copsychus saularis* | Oriental Magpie-robin | Publication and Questionnaire | 5 |
| *Coturnix coturnix* | Common Quail | Questionnaire | 5 |
| *Dicrurus macrocercus* | Black Drongo | Questionnaire | 5 |
| *Eudynamys scolopaceus* | Western Koel | Questionnaire | 5 |
| *Fringilla coelebs* | Common Chaffinch | Questionnaire | 5 |

| | | | |
|---|---|---|---|
| *Garrulax leucolophus* | White-crested Laughingthrush | Publication | 5 |
| *Gubernatrix cristata* | Yellow Cardinal | Questionnaire | 5 |
| *Gyps coprotheres* | Cape Vulture | Publication and Questionnaire | 5 |
| *Halcyon cyanoventris* | Javan Kingfisher | Questionnaire | 5 |
| *Ictinaetus malaiensis* | Black Eagle | Questionnaire | 5 |
| *Ketupa ketupu* | Buffy Fish-owl | Questionnaire | 5 |
| *Leucopsar rothschildi* | Bali Myna | Publication and Questionnaire | 5 |
| *Linaria cannabina* | Common Linnet | Questionnaire | 5 |
| *Lonchura punctulata* | Scaly-breasted Munia | Questionnaire | 5 |
| *Mino dumontii* | Yellow-faced Myna | Questionnaire | 5 |
| *Phasianus colchicus* | Common Pheasant | Questionnaire | 5 |
| *Phodilus badius* | Oriental Bay-owl | Questionnaire | 5 |
| *Pycnonotus zeylanicus* | Straw-headed Bulbul | Publication and Questionnaire | 5 |
| *Serinus serinus* | European Serin | Questionnaire | 5 |
| *Sporophila angolensis* | Chestnut-bellied Seed-Finch | Questionnaire | 5 |
| *Sporophila caerulescens* | Double-collared Seedeater | Questionnaire | 5 |
| *Taeniopygia guttata* | Timor Zebra Finch | Questionnaire | 5 |
| *Trichoglossus ornatus* | Ornate Lorikeet | Questionnaire | 5 |
| *Zonotrichia capensis* | Rufous-collared Sparrow | Questionnaire | 5 |
| *Accipiter gularis* | Japanese Sparrowhawk | Questionnaire | 4 |
| *Accipiter soloensis* | Chinese Sparrowhawk | Questionnaire | 4 |
| *Accipiter trivirgatus* | Crested Goshawk | Questionnaire | 4 |
| *Aplonis panayensis* | Asian Glossy Starling | Questionnaire | 4 |
| *Aprosmictus jonquillaceus* | Jonquil Parrot | Questionnaire | 4 |
| *Aythya nyroca* | Ferruginous Duck | Questionnaire | 4 |
| *Bucorvus leadbeateri* | Southern Ground-hornbill | Questionnaire | 4 |
| *Carduelis carduelis* | European Goldfinch | Publication and Questionnaire | 4 |
| *Chloris chloris* | European Greenfinch | Questionnaire | 4 |
| *Chloropsis cochinchinensis* | Javan Leafbird | Questionnaire | 4 |
| *Chloropsis sonnerati* | Greater Green Leafbird | Publication and Questionnaire | 4 |
| *Coccothraustes coccothraustes* | Hawfinch | Questionnaire | 4 |

| | | | |
|---|---|---|---|
| *Dendrocygna arcuata* | Wandering Whistling-duck | Publication and Questionnaire | 4 |
| *Emberiza aureola* | Yellow-breasted Bunting | Publication and Questionnaire | 4 |
| *Eos histrio* | Red-and-blue Lory | Questionnaire | 4 |
| *Eurystomus orientalis* | Oriental Dollarbird | Questionnaire | 4 |
| *Ficedula mugimaki* | Mugimaki Flycatcher | Questionnaire | 4 |
| *Fulica atra* | Common Coot | Questionnaire | 4 |
| *Geopelia striata* | Zebra Dove | Questionnaire | 4 |
| *Icthyophaga ichthyaetus* | Grey-headed Fish-eagle | Questionnaire | 4 |
| *Kittacincla malabarica* | White-rumped Shama | Publication and Questionnaire | 4 |
| *Macheiramphus alcinus* | Bat Hawk | Questionnaire | 4 |
| *Paroaria dominicana* | Red-cowled Cardinal | Questionnaire | 4 |
| *Pitta elegans* | Elegant Pitta | Questionnaire | 4 |
| *Platylophus galericulatus* | Crested Jay | Questionnaire | 4 |
| *Pomatorhinus montanus* | Chestnut-backed Scimitar-babbler | Questionnaire | 4 |
| *Psittacus timneh* | Timneh Parrot | Publication and Questionnaire | 4 |
| *Pycnonotus aurigaster* | Sooty-headed Bulbul | Questionnaire | 4 |
| *Pycnonotus goiavier* | Yellow-vented Bulbul | Publication and Questionnaire | 4 |
| *Pycnonotus jocosus* | Red-whiskered Bulbul | Publication and Questionnaire | 4 |
| *Pyrrhula pyrrhula* | Eurasian Bullfinch | Questionnaire | 4 |
| *Saltator aurantiirostris* | Golden-billed Saltator | Questionnaire | 4 |
| *Saltator similis* | Green-winged Saltator | Questionnaire | 4 |
| *Saxicola caprata* | Pied Bushchat | Questionnaire | 4 |
| *Sicalis flaveola* | Saffron Finch | Questionnaire | 4 |
| *Spinus atratus* | Black Siskin | Questionnaire | 4 |
| *Spinus magellanicus* | Hooded Siskin | Questionnaire | 4 |
| *Spinus spinus* | Eurasian Siskin | Questionnaire | 4 |
| *Sporophila maximiliani* | Great-billed Seed-finch | Questionnaire | 4 |
| *Tanygnathus lucionensis* | Blue-naped Parrot | Publication | 4 |
| *Trichoglossus euteles* | Olive-headed Lorikeet | Questionnaire | 4 |
| *Acridotheres javanicus* | Javan Myna | Questionnaire | 3 |
| *Acridotheres melanopterus* | Black-winged Myna | Publication and Questionnaire | 3 |

| | | | |
|---|---|---|---|
| *Aegithina tiphia* | Common Iora | Questionnaire | 3 |
| *Alophoixus bres* | Brown-cheeked Bulbul | Questionnaire | 3 |
| *Aythya ferina* | Common Pochard | Questionnaire | 3 |
| *Basilornis celebensis* | Sulawesi Myna | Questionnaire | 3 |
| *Branta ruficollis* | Red-breasted Goose | Questionnaire | 3 |
| *Bubo sumatranus* | Barred Eagle-owl | Questionnaire | 3 |
| *Bycanistes fistulator* | Western Piping Hornbill | Questionnaire | 3 |
| *Ceratogymna atrata* | Black-casqued Hornbill | Questionnaire | 3 |
| *Ceratogymna elata* | Yellow-casqued Hornbill | Questionnaire | 3 |
| *Chloropsis venusta* | Blue-masked Leafbird | Questionnaire | 3 |
| *Corvus macrorhynchos* | Large-billed Crow | Questionnaire | 3 |
| *Cracticus cassicus* | Hooded Butcherbird | Questionnaire | 3 |
| *Cyanoptila cyanomelana* | Blue-and-white Flycatcher | Questionnaire | 3 |
| *Emberiza calandra* | Corn Bunting | Questionnaire | 3 |
| *Emberiza citrinella* | Yellowhammer | Questionnaire | 3 |
| *Falco moluccensis* | Spotted Kestrel | Questionnaire | 3 |
| *Garrulax lugubris* | Black Laughingthrush | Questionnaire | 3 |
| *Geokichla citrina* | Orange-headed Thrush | Publication and Questionnaire | 3 |
| *Gracula robusta* | Nias Hill Myna | Questionnaire | 3 |
| *Horizocerus albocristatus* | Western Long-tailed Hornbill | Questionnaire | 3 |
| *Hydrornis guajanus* | Javan Banded Pitta | Questionnaire | 3 |
| *Lanius schach* | Long-tailed Shrike | Questionnaire | 3 |
| *Leptocoma brasiliana* | Maroon-bellied Sunbird | Questionnaire | 3 |
| *Lonchura maja* | White-headed Munia | Questionnaire | 3 |
| *Lophoceros nasutus* | African Grey Hornbill | Questionnaire | 3 |
| *Lophotriorchis kienerii* | Rufous-bellied Eagle | Questionnaire | 3 |
| *Loriculus pusillus* | Yellow-throated Hanging-parrot | Questionnaire | 3 |
| *Lymnocryptes minimus* | Jack Snipe | Questionnaire | 3 |
| *Mirafra javanica* | Horsfield's Bushlark | Questionnaire | 3 |
| *Ninox scutulata* | Brown Boobook | Questionnaire | 3 |
| *Nisaetus bartelsi* | Javan Hawk-eagle | Publication and Questionnaire | 3 |

| | | | |
|---|---|---|---|
| *Nisaetus lanceolatus* | Sulawesi Hawk-eagle | Questionnaire | 3 |
| *Pernis ptilorhynchus* | Oriental Honey-buzzard | Questionnaire | 3 |
| *Phonygammus keraudrenii* | Trumpet Manucode | Questionnaire | 3 |
| *Poicephalus robustus* | Cape Parrot | Publication | 3 |
| *Pycnonotus bimaculatus* | Orange-spotted Bulbul | Publication and Questionnaire | 3 |
| *Rubigula dispar* | Ruby-throated Bulbul | Questionnaire | 3 |
| *Scissirostrum dubium* | Grosbeak Starling | Questionnaire | 3 |
| *Streptocitta albicollis* | Southern White-necked Myna | Questionnaire | 3 |
| *Tetrao urogallus* | Western Capercaillie | Questionnaire | 3 |
| *Turacoena modesta* | Black Cuckoo-dove | Questionnaire | 3 |
| *Zosterops flavus* | Javan White-eye | Questionnaire | 3 |
| *Acridotheres tertius* | Grey-rumped Myna | Questionnaire | 2 |
| *Acridotheres tricolor* | Grey-backed Myna | Publication and Questionnaire | 2 |
| *Aerodramus fuciphagus* | Edible-nest Swiftlet | Questionnaire | 2 |
| *Alophoixus tephrogenys* | Grey-cheeked Bulbul | Questionnaire | 2 |
| *Amandava formosa* | Green Avadavat | Questionnaire | 2 |
| *Arborophila javanica* | Chestnut-bellied Partridge | Publication | 2 |
| *Artamus leucoryn* | White-breasted Woodswallow | Questionnaire | 2 |
| *Aviceda leuphotes* | Black Baza | Questionnaire | 2 |
| *Brachypteryx montana* | Javan Shortwing | Questionnaire | 2 |
| *Brotogeris pyrrhoptera* | Grey-cheeked Parakeet | Publication and Questionnaire | 2 |
| *Bycanistes brevis* | Silvery-cheeked Hornbill | Questionnaire | 2 |
| *Bycanistes bucinator* | Trumpeter Hornbill | Questionnaire | 2 |
| *Bycanistes subcylindricus* | Grey-cheeked Hornbill | Questionnaire | 2 |
| *Chloropsis media* | Sumatran Leafbird | Questionnaire | 2 |
| *Corvus typicus* | Piping Crow | Questionnaire | 2 |
| *Cyanoloxia brissonii* | Ultramarine Grosbeak | Questionnaire | 2 |
| *Cyornis banyumas* | Hill Blue-flycatcher | Questionnaire | 2 |
| *Cyornis rufigastra* | Mangrove Blue-flycatcher | Questionnaire | 2 |
| *Ducula luctuosa* | White Imperial-pigeon | Questionnaire | 2 |
| *Emberiza cirlus* | Cirl Bunting | Questionnaire | 2 |

| | | | |
|---|---|---|---|
| *Enodes erythrophris* | Fiery-browed Starling | Questionnaire | 2 |
| *Eumyias indigo* | Indigo Flycatcher | Questionnaire | 2 |
| *Ficedula westermanni* | Little Pied Flycatcher | Questionnaire | 2 |
| *Gallinago gallinago* | Common Snipe | Questionnaire | 2 |
| *Gallinago media* | Great Snipe | Questionnaire | 2 |
| *Garrulax bicolor* | Sumatran Laughingthrush | Publication and Questionnaire | 2 |
| *Garrulax mitratus* | Chestnut-capped Laughingthrush | Publication and Questionnaire | 2 |
| *Geokichla dohertyi* | Chestnut-backed Thrush | Questionnaire | 2 |
| *Geokichla interpres* | Chestnut-capped Thrush | Questionnaire | 2 |
| *Geopelia maugeus* | Barred Dove | Questionnaire | 2 |
| *Heleia javanica* | Javan Grey-throated White-eye | Questionnaire | 2 |
| *Lichmera indistincta* | Brown Honeyeater | Questionnaire | 2 |
| *Lonchura fuscata* | Timor Sparrow | Questionnaire | 2 |
| *Lonchura leucogastroides* | Javan Munia | Questionnaire | 2 |
| *Lophoceros alboterminatus* | Crowned Hornbill | Questionnaire | 2 |
| *Lophoceros fasciatus* | Congo Pied Hornbill | Questionnaire | 2 |
| *Loriculus stigmatus* | Sulawesi Hanging-parrot | Questionnaire | 2 |
| *Microhierax fringillarius* | Black-thighed Falconet | Questionnaire | 2 |
| *Mixornis bornensis* | Bold-striped Tit-babbler | Questionnaire | 2 |
| *Oriolus cruentus* | Javan Oriole | Questionnaire | 2 |
| *Orthotomus sepium* | Olive-backed Tailorbird | Questionnaire | 2 |
| *Otus lempiji* | Sunda Scops-owl | Questionnaire | 2 |
| *Philemon buceroides* | Helmeted Friarbird | Questionnaire | 2 |
| *Prinia familiaris* | Bar-winged Prinia | Questionnaire | 2 |
| *Ptilinopus cinctus* | Black-backed Fruit-dove | Questionnaire | 2 |
| *Pycnonotus snouckaerti* | Aceh Bulbul | Questionnaire | 2 |
| *Scolopax rusticola* | Eurasian Woodcock | Questionnaire | 2 |
| *Sphecotheres viridis* | Timor Figbird | Questionnaire | 2 |
| *Sporophila collaris* | Rusty-collared Seedeater | Questionnaire | 2 |
| *Tockus erythrorhynchus* | Red-billed Hornbill | Questionnaire | 2 |
| *Tockus flavirostris* | Eastern Yellow-billed Hornbill | Questionnaire | 2 |

| | | | |
|---|---|---|---|
| *Treron griseicauda* | Grey-cheeked Green-pigeon | Questionnaire | 2 |
| *Trichoglossus weberi* | Flores Lorikeet | Questionnaire | 2 |
| *Trochalopteron yersini* | Collared Laughingthrush | Questionnaire | 2 |
| *Turdus obscurus* | Eyebrowed Thrush | Publication and Questionnaire | 2 |
| *Xanthopsar flavus* | Saffron-cowled Blackbird | Questionnaire | 2 |
| *Zosterops atricapilla* | Black-capped White-eye | Questionnaire | 2 |
| *Zosterops chloris* | Lemon-bellied White-eye | Questionnaire | 2 |
| *Zosterops melanurus* | Sangkar White-eye | Questionnaire | 2 |
| *Alcippe pyrrhoptera* | Javan Fulvetta | Questionnaire | 1 |
| *Arborophila orientalis* | White-faced Partridge | Publication and Questionnaire | 1 |
| *Bycanistes albotibialis* | White-thighed Hornbill | Questionnaire | 1 |
| *Chalcoparia singalensis* | Ruby-cheeked Sunbird | Questionnaire | 1 |
| *Cissa thalassina* | Javan Green Magpie | Publication and Questionnaire | 1 |
| *Cochoa azurea* | Javan Cochoa | Questionnaire | 1 |
| *Cyornis caerulatus* | Large-billed Blue-flycatcher | Questionnaire | 1 |
| *Cyornis unicolor* | Pale Blue-flycatcher | Questionnaire | 1 |
| *Dicaeum maugei* | Red-chested Flowerpecker | Questionnaire | 1 |
| *Dicrurus densus* | Wallacean Drongo | Questionnaire | 1 |
| *Garrulax courtoisi* | Blue-crowned Laughingthrush | Questionnaire | 1 |
| *Garrulax palliatus* | Sunda Laughingthrush | Publication and Questionnaire | 1 |
| *Garrulax rufifrons* | Rufous-fronted Laughingthrush | Publication and Questionnaire | 1 |
| *Geokichla peronii* | Orange-banded Thrush | Questionnaire | 1 |
| *Gerygone sulphurea* | Golden-bellied Gerygone | Questionnaire | 1 |
| *Glaucidium castanopterum* | Javan Owlet | Questionnaire | 1 |
| *Gracula venerata* | Tenggara Hill Myna | Publication and Questionnaire | 1 |
| *Gracupica jalla* | Javan Pied Starling | Publication and Questionnaire | 1 |
| *Harpactes oreskios* | Orange-breasted Trogon | Questionnaire | 1 |
| *Heleia wallacei* | Yellow-spectacled White-eye | Questionnaire | 1 |
| *Laniellus albonotatus* | Spotted Crocias | Publication and Questionnaire | 1 |
| *Lanius cristatus* | Brown Shrike | Questionnaire | 1 |
| *Leiothrix laurinae* | Sumatran Mesia | Questionnaire | 1 |

| | | | |
|---|---|---|---|
| *Loriculus exilis* | Pygmy Hanging-parrot | Questionnaire | 1 |
| *Melloria quoyi* | Black Butcherbird | Questionnaire | 1 |
| *Otus jolandae* | Rinjani Scops-owl | Publication and Questionnaire | 1 |
| *Pachycephala nudigula* | Bare-throated Whistler | Questionnaire | 1 |
| *Philemon inornatus* | Timor Friarbird | Questionnaire | 1 |
| *Pitohui kirhocephalus* | Northern Variable Pitohui | Questionnaire | 1 |
| *Prinia inornata* | Plain Prinia | Questionnaire | 1 |
| *Rhipidura euryura* | White-bellied Fantail | Questionnaire | 1 |
| *Stachyris thoracica* | White-bibbed Babbler | Questionnaire | 1 |
| *Tanygnathus sumatranus* | Azure-rumped Parrot | Questionnaire | 1 |
| *Tockus deckeni* | Von der Decken's Hornbill | Questionnaire | 1 |
| *Zosterops anomalus* | Black-ringed White-eye | Questionnaire | 1 |
| *Zosterops simplex* | Swinhoe's White-eye | Questionnaire | 1 |
| *Zosterops somadikartai* | Togian White-eye | Questionnaire | 1 |
| *Aegithina viridissima* | Green Iora | Questionnaire | 0 |
| *Aethopyga duyvenbodei* | Elegant Sunbird | Questionnaire | 0 |
| *Basilornis galeatus* | Helmeted Myna | Questionnaire | 0 |
| *Larus michahellis* | Yellow-legged Gull | Questionnaire | 0 |
| *Loriculus sclateri* | Sula Hanging-parrot | Questionnaire | 0 |
| *Sporophila cinnamomea* | Chestnut Seedeater | Questionnaire | 0 |
| *Sporophila palustris* | Marsh Seedeater | Questionnaire | 0 |
| *Trochalopteron ngoclinhense* | Golden-winged Laughingthrush | Questionnaire | 0 |
| *Turacoena manadensis* | White-faced Cuckoo-dove | Questionnaire | 0 |

**Table S3.** List of publications used to compile the dataset on birds in markets and other sources

Abi-Said M, R., Outa N, T., Makhlouf H, Amr Z, S., Eid E. 2018. Illegal Trade in Wildlife Species in Beirut, Lebanon. Vertebrate Zoology 68:1-4.

Ahmed A. 2010. Imperilled Custodians of the Night: A study on illegal trade, trapping and utilization of owls in India, New Delhi, India.

Aloufi A, Eid E. 2014. Conservation Perspectives of Illegal Animal Trade at Markets in Tabuk, Saudi Arabia. TRAFFIC Bulletin 26.

Al-Sirhan AR, Al-Bathali O. 2010. Raptor trade in Kuwait bird market. Wildlife Middle East News.

Alves RRN, De Farias Lima JR, Araujo HFP. 2013. The live bird trade in Brazil and its conservation implications: an overview. Bird Conservation International 23:53-65.

Alves RRN, Nogueira EEG, Araujo HFP, Brooks SE. 2010. Bird-keeping in the Caatinga, NE Brazil. Human Ecology 38:147–156.

Alves RRN, Rosa IL. 2010. Trade of Animals Used in Brazilian Traditional Medicine: Trends and Implications for Conservation. Human Ecology 38:691-704.

Angguni T, Mulyani YA, Mardiastuti A. 2021. Bird species contested at songbird competition in Jabodetabek Region, Indonesia. IOP Conference Series: Earth and Environmental Science.

Armstrong OH, Chng SCL. 2020. Distancing the flock: Bird singing competitions fly online to avoid Covid-19. TRAFFIC Bulletin 32:49-55.

Banjade M, Adhikari P, Oh H-S. 2020. Illegal wildlife trade in local markets of Feuang and Mad districts of Vientiane Province, Lao People’s Democratic Republic. Journal of Asia-Pacific Biodiversity 13:511-517.

Brooks-Moizer F, Roberton SI, Edmunds K, Bell D. 2009. Avian Influenza H5N1 and the Wild Bird Trade in Hanoi, Vietnam. Ecology and Society 14.

Buij R, Nikolaus G, Whytock R, Ingram DJ, Ogada D. 2016. Trade of threatened vultures and other raptors for fetish and bushmeat in West and Central Africa. Oryx 50:606-616.

Burivalova Z, Lee TM, Hua F, Lee JSH, Prawiradilaga DM, Wilcove DS. 2017. Understanding consumer preferences and demography in order to reduce the domestic trade in wild-caught birds. Biological Conservation 209:423-431.

Canlas CP, Sy EY, Chng S. 2017. A rapid survey of online trade in live birds and reptiles in the Philippines. TRAFFIC Bulletin 29:58-63.

Chiok WX, Chng SCL. 2021. Trading Faces: Live bird trade on Facebook in Singapore., Southeast Asia Regional Office, Petaling Jaya, Selangor, Malaysia.

Chng SCL, Eaton JA, Krishnasamy K, Shepherd CR, Nijman V. 2015. In the Market for Extinction: An inventory of Jakarta’s bird markets., Petaling Jaya, Selangor, Malaysia.

Chng SCL, Eaton JA. 2016a. In the Market for Extinction: Eastern and Central Java. Petaling Jaya, Selangor, Malaysia.

Chng SCL, Eaton JA. 2016b. Snapshot of an on-going trade: an inventory of birds for sale in Chatuchak weekendmarket, Bangkok, Thailand BirdingASIA 25:24–29.

Chng SCL, Guciano M, Eaton JA. 2016. In the market for extinction: Sukahaji, Bandung, Java, Indonesia. BirdingASIA 26.

Chng SCL, Krishnasamy K, Eaton JA. 2018a. In the market for extinction: the cage bird trade in Bali. FORKTAIL 34:35-41.

Chng SCL, Shepherd CR, Eaton JA. 2018b. In the market for extinction: birds for sale at selected outlets in Sumatra. TRAFFIC Bulletin 30:15-22.

Cottee-Jones HEW, Mittermeier JC, Purba EC, Ashuri NM, Hesdianti E. 2014. An assessment of the parrot trade on Obi Island (North Moluccas) reveals heavy exploitation of the Vulnerable Chattering Lory Lorius garrulus Kukila 18:1-9.

Crook V, Van der Henst E. 2020. Stop wildlife cybercrime in the EU: Online trade in reptiles and birds in Belgium and the Netherlands.

Dai C, Zhang C. 2017. The local bird trade and its conservation impacts in the city of Guiyang, Southwest China. Regional Environmental Change 17:1763-1773.

Dangol BR. 2015. Illegal Wildlife Trade in Nepal: A Case Study from Kathmandu Valley Department of International Environment and Development Studies (Noragric). Norwegian University of Life Sciences (NMBU), Ås, Norway.

Dauphine N. 2008. NOTES ON THE LIVE BIRD TRADE IN NORTHERN PERU. Proceedings of the Fourth International Partners in Flight Conference: Tundra to Tropics:220–222.

Daut EF, Brightsmith DJ, Mendoza AP, Puhakka L, Peterson MJ. 2015. Illegal domestic bird trade and the role of export quotas in Peru. Journal for Nature Conservation 27:44-53.

Davies A, Nuno ANA, Hinsley AMY, Martin RO. 2022. Live wild bird exports from West Africa: insights into recent trade from monitoring social media. Bird Conservation International:1-14.

de Oliveira ES, de Freitas Torres D, Alves RRN. 2020. Wild animals seized in a state in Northeast Brazil: Where do they come from and where do they go? Environment, Development and Sustainability 22:2343–2363.

de Oliveira W, M. Borges AK, Lopes S, Vasconcellos A, Alves R. 2020. Illegal trade of songbirds: an analysis of the activity in an area of northeast Brazil. Journal of Ethnobiology and Ethnomedicine 16.

Desenne P, Strahl SD. 1991. Trade and the conservation status of the family Psittacidae in Venezuela. Bird Conservation International 1:153-169.

Destro GFG, Andrade AFAd, Fernandes Vd, Terribile LC, De Marco P. 2020. Climate suitability as indicative of invasion potential for the most seized bird species in Brazil. Journal for Nature Conservation 58:125890.

do Nascimento CAR, Czaban RE, Alves RRN. 2015. Trends in Illegal Trade of Wild Birds in Amazonas State, Brazil. Tropical Conservation Science 8:1098-1113.

Eaton JA, Leupen BTC, Krishnasamy K. 2017a. Songsters of Singapore: An Overview of the Bird Species in Singapore Pet Shops., Petaling Jaya,  Selangor, Malaysia. .

Eaton JA, Nguyen MDT, Willemsen M, Lee J, Chng SCL. 2017b. Caged in the city: An inventory of birds for sale in Ha Noi and Ho Chi Minh City, Viet Nam. . Southeast Asia Regional Office, Petaling Jaya, Selangor, Malaysia. .

Edmunds K, Roberton SI, Few R, Mahood S, Bui PL, Hunter PR, Bell DJ. 2011. Investigating Vietnam's Ornamental Bird Trade: Implications for Transmission of Zoonoses. EcoHealth 8:63-75.

Eid E, Al Hasani I, Al Share T, Abed O, Amr Z. 2011. Animal Trade in Amman Local Market, Jordan. Jordan Journal of Biological Sciences 4:101 - 108.

Espinosa S, Branch LC, Cueva R. 2014. Road Development and the Geography of Hunting by an Amazonian Indigenous Group: Consequences for Wildlife Conservation. PLoS ONE 9:1-21.

Fa JE, Seymour S, Dupain J, Amin R, Albrechtsen L, Macdonald D. 2006. Getting to grips with the magnitude of exploitation: Bushmeat in the Cross–Sanaga rivers region, Nigeria and Cameroon. Biological Conservation 129:497-510.

Fiennes S, Zhang M, Sun F, Lee TM. 2021. Understanding retail dynamics of a regionally important domestic bird market in Guangzhou, China. Conservation Science and Practice n/a:e487.

Fink C, Toivonen T, Correia RA, Di Minin E. 2021. Mapping the online songbird trade in Indonesia. SocArXiv.

GastaÑAga M, Macleod R, Hennessey B, NÚÑEz JU, Puse E, Arrascue A, Hoyos J, Chambi WM, Vasquez J, Engblom G. 2011. A study of the parrot trade in Peru and the potential importance of internal trade for threatened species. Bird Conservation International 21:76-85.

Gilbert M, Sokha C, Joyner PH, Thomson RL, Poole C. 2012. Characterizing the trade of wild birds for merit release in Phnom Penh, Cambodia and associated risks to health and ecology. Biological Conservation 153:10-16.

Gomes Destro GF, Pimentel TL, Sabaini RM, Borges RC, Barreto R. 2012. Efforts to Combat Wild Animals Trafficking in Brazil in Lameed GA, editor. Biodiversity Enrichment in a Diverse World. InTech, Croatia.

González JA. 2003. Harvesting, local trade, and conservation of parrots in the Northeastern Peruvian Amazon. Biological Conservation 114:437-446.

Greatorex ZF, et al. 2016. Wildlife Trade and Human Health in Lao PDR: An Assessment of the Zoonotic Disease Risk in Markets. PLOS ONE 11:e0150666.

Gunawan., Paridi A, Noske RA. 2017. The illegal trade of Indonesian raptors through social media. Kukila 20:1-11.

Handal EN, Amr ZS, Basha WS, Qumsiyeh MB. 2021. Illegal trade in wildlife vertebrate species in the West Bank, Palestine. Journal of Asia-Pacific Biodiversity 14:636-639.

Herrera M, Hennessey B. 2007. Quantifying the illegal parrot trade in Santa Cruz de la Sierra, Bolivia, with emphasis on threatened species. Bird Conservation International 17:295-300.

Herrera M, Hennessey B. 2008. Monitoring Results of the Illegal Parrot Trade in the Los Pozos Market, Santa Cruz de la Sierra, Bolivia. Proceedings of the Fourth International Partners in Flight Conference: Tundra to Tropics 232-234.

Hung LM, Craik RC. 2016. Notes on the trading of some threatened and endemic species from Vietnam BirdingASIA 26:17–21.

Hussain A, Khan AA. 2021. Wild birds trade in Dera Ismael Khan and Bannu divisions of Khyber PakhtunKhwa (KPK) Province, Pakistan. Brazilian Journal of Biology 83:1-7.

Indraswari K, Friedman RS, Noske R, Shepherd CR, Biggs D, Susilawati C, Wilson C. 2020. It's in the news: Characterising Indonesia's wild bird trade network from media-reported seizure incidents. Biological Conservation 243:108431.

Iqbal M. 2016. Predators become prey! Can Indonesian raptors survive online bird trading? BirdingASIA 25:30–35.

Iskandar BS, Iskandar J, Partasasmita R. 2019. Hobby and business on trading birds: Case study in bird market of Sukahaji, Bandung, West Java and Splendid, Malang, East Java (Indonesia). Biodiversitas 20:1316-1332.

Iskandar J, Iskandar BS, Mulyanto D, Alfian RL, Partasasmita R. 2020. Traditional ecological knowledge of the bird traders on bird species bird naming, and bird market chain: A case study in bird market Pasty Yogyakarta, Indonesia. BIODIVERSITAS 21:2586-2602.

Khaing TT. 2019. Parakeet Trade in Shwesettaw Wildlife Area, Minbu Township, Magway Region, During Pagoda Festival. International Journal of Innovative Science and Research Technology 4:252-255.

Krishna VV, Darras K, Grass I, Mulyani YA, Prawiradilaga DM, Tscharntke T, Qaim M. 2019. Wildlife trade and consumer preference for species rarity: an examination of caged-bird markets in Sumatra. Environment and Development Economics 24:339-360.

Krishnasamy K, Stoner S. 2016. Trading Faces: A Rapid Assessment on the use of Facebook to Trade Wildlife in Peninsular Malaysia. Petaling Jaya, Selangor, Malaysia.

Leupen BTC, Gomez L, Shepherd CR, Nekaris KA-I, Imron MA, Nijman V. 2020. Thirty years of trade data suggests population declines in a once common songbird in Indonesia. European Journal of Wildlife Research 66:1-11.

Leupen BTC, Shepherd L, Shepherd CR, Damianou E, Nijman V. 2022. Market surveys in Mataram, Lombok, illustrate the expanse of legaland illegal Indonesian bird trade networks. InJAST 3:42-52.

Matias CAR, Oliveira VM, Rodrigues DP, Siciliano S. 2012. Summary of the Bird Species Seized in the Illegal Trade in Rio de Janeiro, Brazil   TRAFFIC Bulletin 24:83-86.

Maulany RI, Mutmainnah A, Nasri N, Achmad A, Ngakan PO. 2021. Tracing Current Wildlife Trade: An Initial Investigation in Makassar City, Indonesia. Forest and Society 5:277-287.

McKean S, Mander M, Diederichs N, Ntuli L, Mavundla K, Williams V, Wakelin J. 2013. The impact of traditional use on vultures in South Africa. Vulture News 65:15-36.

Nguyen M, Willemsen M. 2016. A rapid assessment of e-commerce wildlife trade in Viet Nam. TRAFFIC Bulletin 28:53-55.

Nijman V, Campera M, Imron MA, Ardiansyah A, Langgeng A, Dewi T, Hedger K, Hendrik R, Nekaris KA. 2021c. The Role of the Songbird Trade as an Anthropogenic Vector in the Spread of Invasive Non-Native Mynas in Indonesia. Life 11.

Nijman V, Morcatty TQ, Feddema K, Campera M, Nekaris KAI. 2022. Disentangling the Legal and IllegalWildlife Trade–Insights from Indonesian Wildlife Market Surveys Animals 12:1-21.

Nijman V, Sari SL, Siriwat P, Sigaud M, Nekaris KA-I. 2017. Records of four Critically Endangered songbirds in the markets of Java suggest domestic trade is a major impediment to their conservation. BirdingASIA 27:20-25.

Nikolaus G. 2001. Bird exploitation for traditional medicine in Nigeria. Malimbus 23:45-55.

Pangau-Adam M, Noske R. 2010. Wildlife Hunting and Bird Trade in Northern Papua (Irian Jaya), Indonesia. Pages 73-85. Ethno.

Petrozzi F. Bushmeat and fetish trade of birds in West Africa: A review. Life and environment 68:51-64.

Phassaraudomsak M, Krishnasamy K, Chng SCL. 2019. Trading Faces: Online trade of Helmeted and other hornbill species on Facebook in Thailand. TRAFFIC, Southeast Asia Regional Office, Petaling Jaya, Malaysia.

Phassaraudomsak M, Krishnasamy K. 2018. Trading Faces: A rapidassessment on the use of Facebook to trade inwildlife in Thailand. Petaling Jaya, Selangor, Malaysia.

ProFauna. 2009. Wildlife Trade Survey on the Bird markets in Java.

Razkallah I, Atoussi S, Telailia S, Abdelghani M, Zihad B, Moussa H. 2019. Illegal wild birds' trade in a street market in the region of Guelma, north-east of Algeria. Avian Biology Research 12:96-102.

Regueira RFS, Bernard E. 2012. Wildlife sinks: Quantifying the impact of illegal bird trade in street markets in Brazil. Biological Conservation 149:16-22.

Reuter KE, Clarke TA, LaFleur M, Rodriquez L, Hanitriniaina S, Schaefer MS. 2017. Trade of parrots in urban areas of Madagascar. MADAGASCAR CONSERVATION & DEVELOPMENT 12:41-48.

RSCN. 2020. The status of illegal trade in wild birds at the local market and online with focus on soaring birds.

Setiyani AD, Ahmadi MA. 2020. An overview of illegal parrot trade in Maluku and North Maluku Provinces. Forest and Society 4:48-60.

Shanee N. 2012. Trends in local wildlife hunting, trade and control in the Tropical Andes Biodiversity Hotspot, northeastern Peru. Endangered Species Research 19:177-186.

Shepherd CR, Eaton JA, Chng SCL. 2015. Pittas for a pittance: observations on the little known illegal trade in Pittidae in west Indonesia BirdingASIA 24:18-20.

Shepherd CR, Eaton JA, Chng SCL. 2016. Nothing to laugh about – the ongoing illegal trade in laughingthrushes (Garrulax species) in the bird markets of Java, Indonesia. Bird Conservation International 26:524-530.

Shepherd CR, Leupen BTC, Siriwat P, Nijman V. 2020a. International wildlife trade, avian influenza, organised crime and the effectiveness of CITES: The Chinese hwamei as a case study. Global Ecology and Conservation 23:e01185.

Shepherd CR, Leupen BTC. 2021. Incidence of protected and illegally sourced birds at birdmarkets in Makassar, Sulawesi. JOURNAL OF ASIAN ORNITHOLOGY 37:29–33.

Shepherd CR. 2006. The bird trade in Medan, north Sumatra: an overview BirdingASIA 5:16-24.

Shepherd CR. 2012. The owl trade in Jakarta, Indonesia: a spot check on the largest bird markets BirdingASIA 18:58–59.

Shivambu TC, Shivambu N, Downs CT. 2022. An assessment of avian species sold in the South African pet trade. African Journal of Ecology n/a.

Siriwat P, Nekaris KAI, Nijman V. 2020. Digital media and the modern-day pet trade: a test of the 'Harry Potter effect' and the owl trade in Thailand. Endangered Species Research 41:7-16.

Siriwat P, Nijman V. 2020. Wildlife trade shifts from brick-and-mortar markets to virtual marketplaces: A case study of birds of prey trade in Thailand. Journal of Asia-Pacific Biodiversity 13:454-461.

Soewu DA, Dedeke GA, Ojo VA, Soewu OK. 2016. Trade in Non-Mammalian Wild Animals for Traditional African Medicine in Ogun State, Nigeria. Global Journals Inc. 16.

Soorae PS, Al Hemeri A, Al Shamsi A, Al Suwaidi K. 2008. A Survey of the Trade in Wildlife as Pets in the United Arab Emirates. TRAFFIC Bulletin 22:41-46.

Su S, Cassey P, Vall-llosera M, Blackburn TM. 2015. Going Cheap: Determinants of Bird Price in the Taiwanese Pet Market. PLOS ONE 10:e0127482.

Tamalene MN, Said H, Kartika K. 2019. Local knowledge and community behavior in the exploitation of parrots in surrounding area of aketajawe lolobata national park. Biosfer: Jurnal Pendidikan Biologi 12.

TRAFFIC. 2021. Calling for Compassion: Countering Vietnam's Songbird Demand with Buddhist Philosophy.

Vall-llosera M, Cassey P. 2017. Physical attractiveness, constraints to the trade and handling requirements drive the variation in species availability in the Australian cagebird trade. Ecological Economics 131:407-413.

Whiting MJ, Williams VL, Hibbitts TJ. 2011. Animals Traded for Traditional Medicine at the Faraday Market in South Africa: Species Diversity and Conservation Implications. Journal of Zoology 284:84–96.

Xayyasith S, Douangboubpha B, Chaiseha Y. 2020. Recent surveys of the bird trade in local markets in central Laos. Forktail 36:47-55.

**Table S4.** The 98 bird species with trade prevalence scores of 5 or above that are not listed in CITES Appendices I or II. Those marked with an asterisk are listed in CITES Appendix III.

| **Common name** | **Scientific name** | **Order** |
|---|---|---|
| Chukar | *Alectoris chukar* | Galliformes |
| Lady Amherst's Pheasant | *Chrysolophus amherstiae* | Galliformes |
| Northern Bobwhite | *Colinus virginianus* | Galliformes |
| Common Quail | *Coturnix coturnix* | Galliformes |
| Great Curassow* | *Crax rubra* | Galliformes |
| Red Junglefowl | *Gallus gallus* | Galliformes |
| Siamese Fireback | *Lophura diardi* | Galliformes |
| Kalij Pheasant* | *Lophura leucomelanos* | Galliformes |
| Silver Pheasant | *Lophura nycthemera* | Galliformes |
| Ocellated Turkey* | *Meleagris ocellata* | Galliformes |
| Helmeted Curassow* | *Pauxi pauxi* | Galliformes |
| Indian Peafowl* | *Pavo cristatus* | Galliformes |
| Crested Guan* | *Penelope purpurascens* | Galliformes |
| Common Pheasant | *Phasianus colchicus* | Galliformes |
| Crested Partridge | *Rollulus rouloul* | Galliformes |
| Satyr Tragopan* | *Tragopan satyra* | Galliformes |
| Egyptian Goose | *Alopochen aegyptiaca* | Anseriformes |
| Northern Pintail | *Anas acuta* | Anseriformes |
| Common Teal | *Anas crecca* | Anseriformes |
| Mallard | *Anas platyrhynchos* | Anseriformes |
| Canada Goose | *Branta canadensis* | Anseriformes |
| Muscovy Duck | *Cairina moschata* | Anseriformes |
| Black Swan | *Cygnus atratus* | Anseriformes |
| Mute Swan | *Cygnus olor* | Anseriformes |
| Black-bellied Whistling-duck* | *Dendrocygna autumnalis* | Anseriformes |
| Fulvous Whistling-duck* | *Dendrocygna bicolor* | Anseriformes |
| White-faced Whistling-duck | *Dendrocygna viduata* | Anseriformes |
| African Pygmy-goose | *Nettapus auritus* | Anseriformes |
| Spur-winged Goose | *Plectropterus gambensis* | Anseriformes |
| Northern Shoveler | *Spatula clypeata* | Anseriformes |
| Garganey | *Spatula querquedula* | Anseriformes |
| Speckled Pigeon | *Columba guinea* | Columbiformes |
| Rock Dove | *Columba livia* | Columbiformes |
| Laughing Dove | *Spilopelia senegalensis* | Columbiformes |
| European Turtle-dove | *Streptopelia turtur* | Columbiformes |
| Mourning Dove | *Zenaida macroura* | Columbiformes |
| Western Koel | *Eudynamys scolopaceus* | Cuculiformes |
| Purple Gallinule | *Porphyrio martinicus* | Gruiformes |

| | | |
|---|---|---|
| Great Blue Turaco | *Corythaeola cristata* | Musophagiformes |
| Violet Turaco | *Musophaga violacea* | Musophagiformes |
| Marabou | *Leptoptilos crumenifer* | Ciconiiformes |
| Lesser Adjutant | *Leptoptilos javanicus* | Ciconiiformes |
| Great White Egret | *Ardea alba* | Pelecaniformes |
| Cattle Egret | *Bubulcus ibis* | Pelecaniformes |
| Little Egret | *Egretta garzetta* | Pelecaniformes |
| Great White Pelican | *Pelecanus onocrotalus* | Pelecaniformes |
| King Vulture* | *Sarcoramphus papa* | Cathartiformes |
| Common Hoopoe | *Upupa epops* | Bucerotiformes |
| Javan Kingfisher | *Halcyon cyanoventris* | Coraciiformes |
| Collared Kingfisher | *Todiramphus chloris* | Coraciiformes |
| Chestnut-eared Araçari* | *Pteroglossus castanotis* | Piciformes |
| Red-breasted Toucan* | *Ramphastos dicolorus* | Piciformes |
| Rose-ringed Parakeet | *Alexandrinus krameri* | Psittaciformes |
| Budgerigar | *Melopsittacus undulatus* | Psittaciformes |
| Common Myna | *Acridotheres tristis* | Passeriformes |
| Red-winged Blackbird | *Agelaius phoeniceus* | Passeriformes |
| Eurasian Skylark | *Alauda arvensis* | Passeriformes |
| Cut-throat Finch | *Amadina fasciata* | Passeriformes |
| Red Avadavat | *Amandava amandava* | Passeriformes |
| Northern Cardinal | *Cardinalis cardinalis* | Passeriformes |
| Amazonian Umbrellabird* | *Cephalopterus ornatus* | Passeriformes |
| Gouldian Finch | *Chloebia gouldiae* | Passeriformes |
| Oriental Magpie-robin | *Copsychus saularis* | Passeriformes |
| Bluethroat | *Cyanecula svecica* | Passeriformes |
| Blue Jay | *Cyanocitta cristata* | Passeriformes |
| Black Drongo | *Dicrurus macrocercus* | Passeriformes |
| European Robin | *Erithacus rubecula* | Passeriformes |
| Common Waxbill | *Estrilda astrild* | Passeriformes |
| Black-rumped Waxbill | *Estrilda troglodytes* | Passeriformes |
| African Silverbill | *Euodice cantans* | Passeriformes |
| Southern Red Bishop | *Euplectes orix* | Passeriformes |
| Common Chaffinch | *Fringilla coelebs* | Passeriformes |
| White-crested Laughingthrush | *Garrulax leucolophus* | Passeriformes |
| Eurasian Jay | *Garrulus glandarius* | Passeriformes |
| House Finch | *Haemorhous mexicanus* | Passeriformes |
| Asian Fairy-bluebird | *Irena puella* | Passeriformes |

| | | |
|---|---|---|
| African Firefinch | *Lagonosticta rubricata* | Passeriformes |
| Golden-breasted Starling | *Lamprotornis regius* | Passeriformes |
| Common Linnet | *Linaria cannabina* | Passeriformes |
| White-rumped Munia | *Lonchura striata* | Passeriformes |
| Northern Mockingbird | *Mimus polyglottos* | Passeriformes |
| Yellow-faced Myna | *Mino dumontii* | Passeriformes |
| House Sparrow | *Passer domesticus* | Passeriformes |
| Blue-winged Pitta | *Pitta moluccensis* | Passeriformes |
| Black-throated Finch | *Poephila cincta* | Passeriformes |
| Island Canary | *Serinus canaria* | Passeriformes |
| European Serin | *Serinus serinus* | Passeriformes |
| Black-and-white Mannikin | *Spermestes bicolor* | Passeriformes |
| Chestnut-bellied Seed-Finch | *Sporophila angolensis* | Passeriformes |
| Double-collared Seedeater | *Sporophila caerulescens* | Passeriformes |
| Lined Seedeater | *Sporophila lineola* | Passeriformes |
| Yellow-bellied Seedeater | *Sporophila nigricollis* | Passeriformes |
| Zebra Finch | *Taeniopygia guttata* | Passeriformes |
| Paradise Tanager | *Tangara chilensis* | Passeriformes |
| Rufous-bellied Thrush | *Turdus rufiventris* | Passeriformes |
| Pin-tailed Whydah | *Vidua macroura* | Passeriformes |
| Blue-black Grassquit | *Volatinia jacarina* | Passeriformes |
| Rufous-collared Sparrow | *Zonotrichia capensis* | Passeriformes |
